# Synergistic Coding of Carbon Dioxide and a Human Sweat Odorant in the Mosquito Brain

**DOI:** 10.1101/2020.11.02.365916

**Authors:** Shruti Shankar, Genevieve M. Tauxe, Emma D. Spikol, Ming Li, Omar S. Akbari, Diego Giraldo, Conor J. McMeniman

**Affiliations:** W. Harry Feinstone Department of Molecular Microbiology and Immunology, Johns Hopkins Malaria Research Institute, Johns Hopkins Bloomberg School of Public Health, Johns Hopkins University, Baltimore, MD 21205, USA; The Solomon H. Snyder Department of Neuroscience, Johns Hopkins University School of Medicine, Baltimore, MD 21205, USA; Section of Cell and Developmental Biology, University of California, San Diego, La Jolla, CA, 92093, USA

## Abstract

The yellow fever mosquito *Aedes aegypti* employs olfaction to locate humans. We applied neural activity mapping to define the molecular and cellular logic of how the mosquito brain is wired to detect two human odorants that are attractive when blended together. We determined that the human breath volatile carbon dioxide (CO_2_) is detected by the largest unit of olfactory coding in the antennal lobe of the mosquito brain. Synergistically, CO_2_ detection gates pre-synaptic calcium signaling in olfactory sensory neuron axon terminals that innervate unique antennal lobe regions tuned to the human sweat odorant L-(+)-lactic acid. We propose that simultaneous detection of the signature human volatiles CO_2_ and L-(+)-lactic acid disinhibits a multimodal olfactory network for hunting humans in the mosquito brain.

## Introduction

Bloodthirsty female *A. aegypti* mosquitoes detect and navigate towards a plethora of physical and chemosensory cues emitted by the human body (Cardé, 2015). Of these cues, human scent is a powerful mosquito attractant, comprising of a complex bouquet of hundreds of volatile chemicals derived from sweat, breath and the human skin microbiome (Dormont et al., 2013). Despite recent advances in our understanding of mosquito chemoreception (Carey et al., 2010; DeGennaro et al., 2013; Lahondère et al., 2020; McMeniman et al., 2014; Raji et al., 2019), how signature components of human scent blend together and are integrated by the mosquito peripheral nervous system and brain to drive behavioral attraction to humans is largely unknown.

L-(+)-lactic acid is a predominant compound found in human sweat that is alone unattractive to *A. aegypti*, yet potently synergizes with CO_2_ exhaled in breath to elicit olfactory attraction when these two stimuli are combined together (Acree et al., 1968; Eiras & Jepson, 1991; McMeniman et al., 2014; Smith et al., 1970). These two odorants are detected at divergent locations in the mosquito olfactory system. L-(+)-lactic acid is detected by specialized olfactory sensory neurons (OSNs) located on the antennae (Davis & Sokolove, 1976), whereas CO_2_ is detected by dedicated OSNs housed on the maxillary palps (Grant et al., 1995).

Peripheral OSN chemoreceptors are required for detection and behavioral synergism between these two olfactory stimuli. The ionotropic receptor (IR) co-receptor *IR8a* putatively forms IR complexes tuned to L-(+)-lactic acid and related acidic volatiles (Raji et al., 2019), while the gustatory receptor (Gr) 1/2/3 CO_2_ receptor complex (McMeniman et al., 2014) detects CO_2_. Previous extracellular recordings from acid-sensitive OSNs on the mosquito antennae (Davis & Sokolove, 1976) indicate that synergy between CO_2_ and L-(+)-lactic acid likely does not occur though peripheral mechanisms operating at dendrites during co-stimulation (Su et al., 2012; Xu et al., 2020). This hints towards integration of these human-related olfactory cues at higher levels of olfactory coding in the mosquito brain.

In most insects, OSNs expressing unique complements of chemoreceptors project from the periphery to spatially defined clusters of synaptic connectivity in the antennal lobe called glomeruli, where odor information is locally encoded and processed. In *Drosophila*, the axonal processes of OSNs expressing unique complements of chemoreceptors project from the peripheral sensory appendages such as the antenna and maxillary palp to stereotypically and spatially defined glomeruli within the antennal lobe (Couto et al., 2005; Vosshall et al., 2000). As a first step towards facilitating in-depth neuroanatomical and functional studies of the *A. aegypti* antennal lobe, we previously determined that this olfactory center contains ~ 80 units of glomerular synaptic connectivity in this mosquito species using neural staining methods (Shankar & McMeniman, 2020).

Here, we report the development of optimized transgenic tools for neuroanatomy and functional imaging of OSNs in *A. aegypti* expressing three major olfactory co-receptor genes: odorant receptor co-receptor (*orco*), ionotropic receptor co-receptor *IR8a*, and the CO_2_ receptor complex subunit *Gr1*. We applied these tools to illuminate synergistic coding of CO_2_ and L-(+)-lactic acid detection in the *A. aegypti* antennal lobe. Using receptor-to-glomerulus labeling, we determined that *Gr1* (+) CO_2_ receptor neurons innervate the largest glomerulus in the *A. aegypti* antennal lobe. In contrast, *IR8a* (+) and *orco* (+) OSNs innervate expanded complements of glomeruli in this brain center. Using neural activity mapping with the photoconvertible neural activity sensor CaMPARI2, we discovered a striking pattern of synergistic odor coding putatively operational between the CO_2_ receptor glomerulus and the axon terminals of *IR8a* (+) OSNs tuned to L-(+)- lactic acid innervating the antennal lobe. Odor-evoked responses in two specific *IR8a* (+) glomeruli were dramatically elevated when mosquitoes were co-stimulated with L-(+)-lactic acid and CO_2_, yet these glomeruli appeared silent when stimulated with either ligand alone. We propose that CO_2_ detection unleashes behavioral attraction towards L-(+)-lactic acid – and possibly other human odorants – via disinhibitory neural circuitry operating at the first olfactory synapse in the *A. aegypti* antennal lobe.

## Results

To provide genetic access to OSNs expressing different classes of chemoreceptors in the *A. aegypti* olfactory system, we first applied CRISPR-Cas9 genome engineering (Kistler et al., 2015) and *Mos1-mariner* transposition (Coates et al., 1998) to integrate components of the *QF2/QUAS* system (Riabinina et al., 2015) into the mosquito genome for binary reporter transgene expression. To generate transgenic *A. aegypti* chemoreceptor-*QF2* driver lines, we used CRISPR-Cas9 mediated homologous recombination to insert a *T2A-QF2* in-frame fusion cassette (Diao & White, 2012; Matthews et al., 2019) into the coding exons of three major olfactory co-receptor genes: *orco, IR8a* and *Gr1* (*Figure 1—figure supplement 1A*).

Using this strategy, *QF2* was integrated in-frame into Exon 3 of each target gene, placing expression of this transcription factor under control of endogenous regulatory elements for each locus. All driver lines included a visible *3xP3-DsRed2* eye marker to facilitate identification of transgenic individuals. Each chemoreceptor-*QF2* driver line (*orco^QF2Red^, IR8a^QF2Red^* and *Gr1^QF2Red^*) was then crossed with a *QUAS-mCD8::GFP* responder strain that we generated, and olfactory tissue of progeny were surveyed for the presence of membrane-tethered green fluorescent protein (GFP) in OSNs (*Figure 1A–C*).

**Figure 1.**
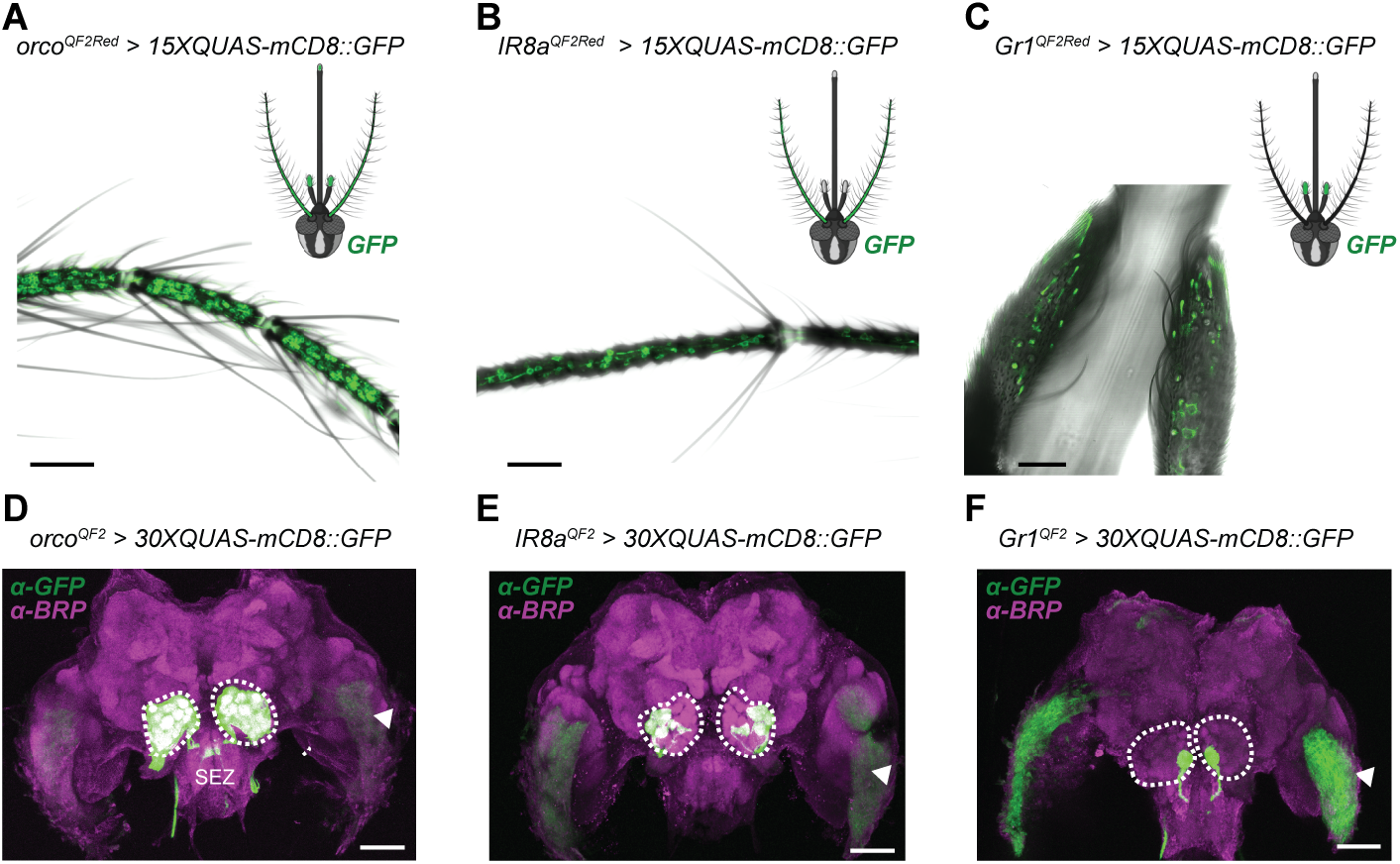
OSNs expressing divergent chemoreceptor gene families project centrally to defined regions of the *Aedes aegypti* antennal lobe. (**A** to **C**) QF2/QUAS system-mediated fluorescent labeling with membrane-tethered GFP (mCD8::GFP) of OSNs expressing (A) *orco*, (B) *IR8a* and (C) *Gr1* co-receptors on *A. aegypti* olfactory appendages. Scale bars: 50 μm. (**D** to **F**) Projection patterns of OSNs expressing these co-receptors into the central mosquito brain. Maximum intensity projections for (D) *orco^QF2^* > *30XQUAS-mCD8::GFP* and (E) *IR8a^QF2^ > 30XQUAS-mCD8::GFP* genotypes, and a single posterior z-slice for the (F) *Gr1^QF2^ > 30XQUAS-mCD8::GFP* genotype are shown at 10X magnification. The antennal lobes are encircled in white. SEZ: subesophageal zone. Arrows indicate expression of the *3xP3-ECFP* marker for the *QUAS* responder transgene in the outer optic lobes.

Confocal analyses of adult female peripheral sensory appendages revealed strong GFP labeling of *orco* (+) OSN dendrites and cell bodies on the antenna, maxillary palp and labella of the proboscis (*Figure 1A* and *Figure 1—figure supplement 2A-E*). Strong labeling of *IR8a* (+) OSNs within the antennal flagellum (*Figure 1B*) and *Gr1* (+) OSNs in maxillary palp tissue (*Figure 1C*) was also detected. Expression patterns from *QF2* knock-ins were consistent with a previous *A. aegypti LVPib12* strain neurotranscriptome analysis (Matthews et al., 2016) that revealed broad *orco* expression across adult olfactory tissues, along with strong expression of *IR8a* and *Gr1* in antennal and maxillary palp tissue, respectively.

We observed that dendrites of *orco* (+) neurons on the mosquito antenna were grossly localized to hair-like trichoid sensilla (*Figure 1A*). Dendrites of *IR8a* (+) neurons appeared confined to grooved-peg sensilla on the antenna (*Figure 1B*). *Gr1* (+) and *orco* (+) neurons were found in capitate peg sensilla on the maxillary palp (*Figure 1C*). These latter two classes of sensilla are the locations for OSN-based detection of L-(+)-lactic acid (Davis & Sokolove, 1976) and CO_2_ (McMeniman et al., 2014), respectively, validating the neuroanatomical specificity of our transgenic labeling approach.

To examine central projection patterns of OSNs expressing these chemoreceptors into the *A. aegypti* brain, we next generated marker-free driver strains (*orco^QF2^, IR8a^QF2^* and *Gr1^QF2^*) devoid of their original fluorescent eye markers. Using *Cre-LoxP* mediated cassette excision, we removed *3xP3* marker cassettes from our original chemoreceptor driver strains (*Figure 1—figure supplement 1B–D*), as they induced spurious labeling in the central brain when integrated at these genomic loci (*Figure 1—figure supplement 3A–I*).

After crossing each marker-free driver strain to our *QUAS-mCD8::GFP* responder strain, we dissected brains of adult female progeny and performed immunohistochemistry analyses with a primary antibody directed against the pre-synaptic protein Bruchpilot (BRP) (Hofbauer et al., 2009) to demarcate glomerular boundaries of neuropils in the antennal lobe, and anti-GFP antibody to amplify mCD8::GFP signal. We then performed confocal imaging and complete 3D morphological reconstructions of antennal lobes with labeled projections from *orco* (+), *IR8a* (+) and *Gr1* (+) OSNs (*Figure 1D–F* and *Figure 2A–C*).

**Figure 2.**
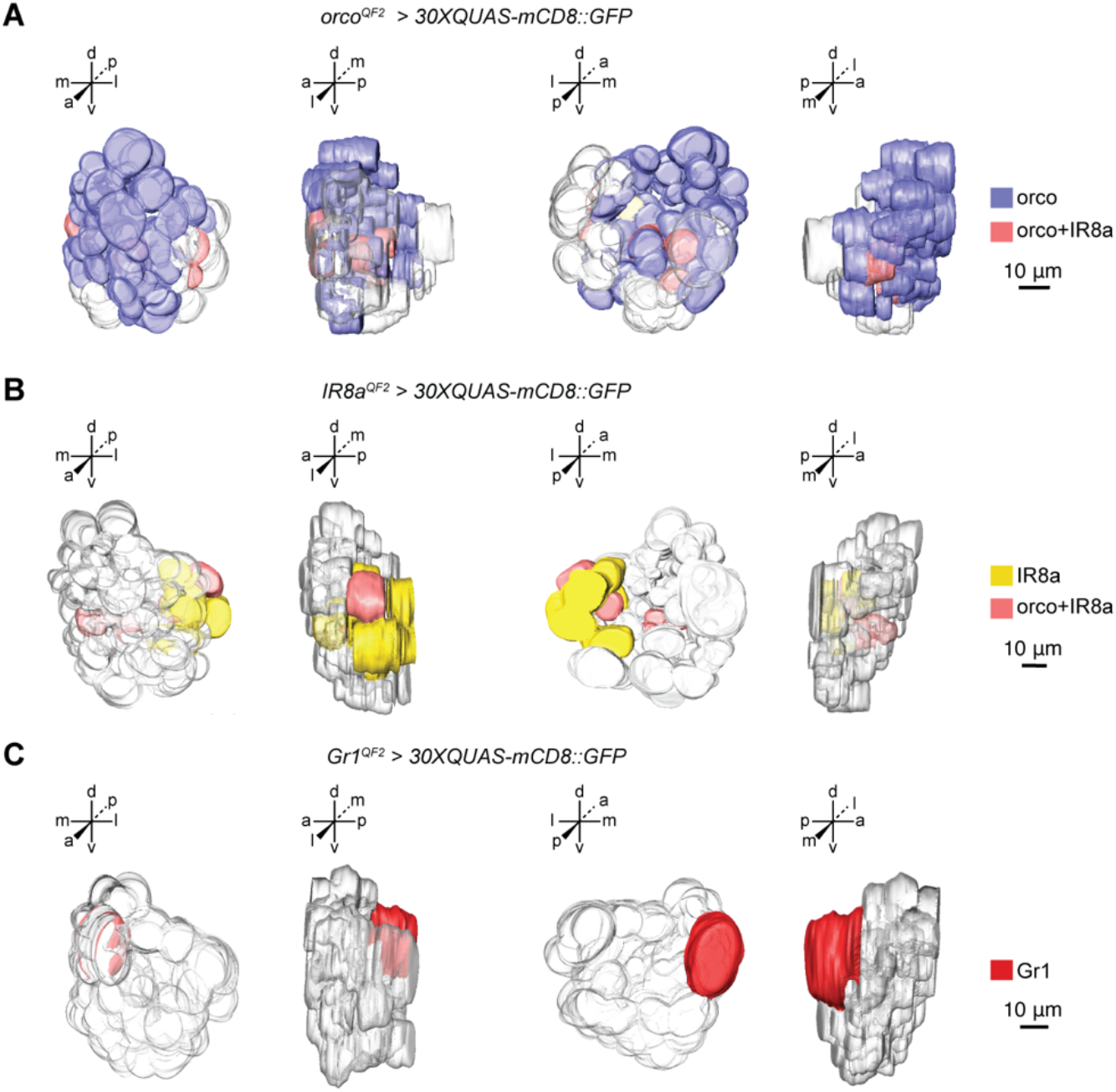
Three-dimensional models of the female *Aedes aegypti* antennal lobe highlighting the innervation patterns of *orco, IR8a* and *Gr1* (+) neurons. **(A)** 3D reconstructed model of *orco^QF2^ > 30XQUAS-mCD8::GFP* left antennal lobe. *orco* (+) glomeruli are colored blue and *orco* (-) glomeruli are transparent **(B)** 3D reconstructed model of *IR8a^QF2^ > 30XQUAS-mCD8::GFP* left antennal lobe. *IR8a* (+) glomeruli are colored yellow and *IR8a* (-) glomeruli are transparent. Glomeruli putatively co-innervated by *orco* and *IR8a* neurons as determined by spatial mapping are colored in pink in A and B. **(C)** 3D reconstructed model of *Gr1^QF2^ > 30XQUAS-mCD8::GFP* left antennal lobe. The *Gr1* (+) glomerulus MD1 is colored red and all other *Gr1* (-) glomerulus are transparent. 3D models are arranged (left to right) in the anterior, lateral, posterior and medial orientations. Scale bar: 10 μm.

Across this dataset, we defined ~ 79 total glomeruli (79 ± 3, mean ± SEM) in each reconstructed antennal lobe (*Figure 2—figure supplement 1A*). This count was consistent with our previous estimate of ~ 80 total glomeruli constituting the female *A. aegypti* antennal lobe based on reconstructions with synaptic staining alone (Shankar & McMeniman, 2020). On average, we counted ~ 63 *orco* (+) glomeruli (63 ± 1, mean ± SEM), 15 *IR8a* (+) glomeruli and 1 *Gr1* (+) glomerulus labeled with GFP in reconstructed antennal lobes from each genotype (*Figure 2—figure supplement 1B*).

Using a systematic reference key for *A. aegypti* antennal lobe nomenclature (Shankar & McMeniman, 2020), we previously determined that 63 out of 80 total glomeruli (~79%) could be found in stereotypical spatial positions in this mosquito species based on synaptic staining alone. In this study with transgenic labeling, cross referencing nc82 and GFP signals markedly improved our ability to define glomerular boundaries and discern the spatial arrangement of glomeruli within the antennal lobe *in vitro*. Indeed, we determined that 74 out of the 79 total glomeruli (~94%) that we annotated could now be assigned to spatially conserved positions across reconstructions (*Figure 2—figure supplement 1C*). Of these spatially invariant glomeruli, we estimate that *orco* (+) OSNs innervate the largest number of glomeruli (56/74) across several spatial regions of the antennal lobe, *IR8a* (+) neurons innervate a sparser subset of glomeruli (15/74) and *Gr1* (+) neurons (1/74) innervate a single, posteriorly positioned glomerulus in the mediodorsal antennal lobe region (*Figure 2—figure supplement 1C–D*).

Using spatial mapping, we estimate a subset of 6 out of these 74 spatially invariant glomeruli are putatively both *orco* (+) and *IR8a* (+) (*Figure 2—figure supplement 1C–D*), indicative of co-expression of these two genes in some OSN types. In addition, 8 of these 74 glomeruli in the ventral region of the antennal lobe appear not to be labeled by any of these chemoreceptor driver lines (*Figure 2—figure supplement 1C–D*). We suggest these latter glomeruli may receive innervations from OSNs expressing other chemoreceptors such as those complexed with IR co-receptors IR25a and IR76b, which are known to project to the antennal lobe in *Drosophila* (Silbering et al., 2011). A minor subset of glomeruli in each reconstructed antennal lobe (6 ± 0.8, mean ± SEM) were also found in variable spatial positions (*Figure 2—figure supplement 1C* and *1E*). Typically, these variant glomeruli were *orco* (+) and sparsely labeled, or unlabeled and situated deep in the antennal lobe making delineation of clear glomerular boundaries in these instances difficult.

As a guide, we assembled representative 2D maps of chemoreceptor innervation in the antennal lobe for *orco* (+) OSNs (*Figure 2—figure supplement 2*), *IR8a* (+) OSNs (*Figure 2—figure supplement 3*) and *Gr1* (+) OSNs (*Figure 2—figure supplement 4*). Additionally, we visualized *orco* (+) neurons innervating the taste center of the insect brain known as the subesophageal zone (SEZ) (*Figure 2—figure supplement 5*), consistent with the projection pattern of *orco* (+) OSNs in the African malaria mosquito *Anopheles gambiae* (Riabinina et al., 2016). Volumetric analysis of *A. aegypti* antennal lobe glomeruli from each of these clusters of chemoreceptor innervation revealed that the *Gr1* (+) glomerulus MD1 is the largest glomerulus in the antennal lobe (*Figure 3A-B*).

**Figure 3.**
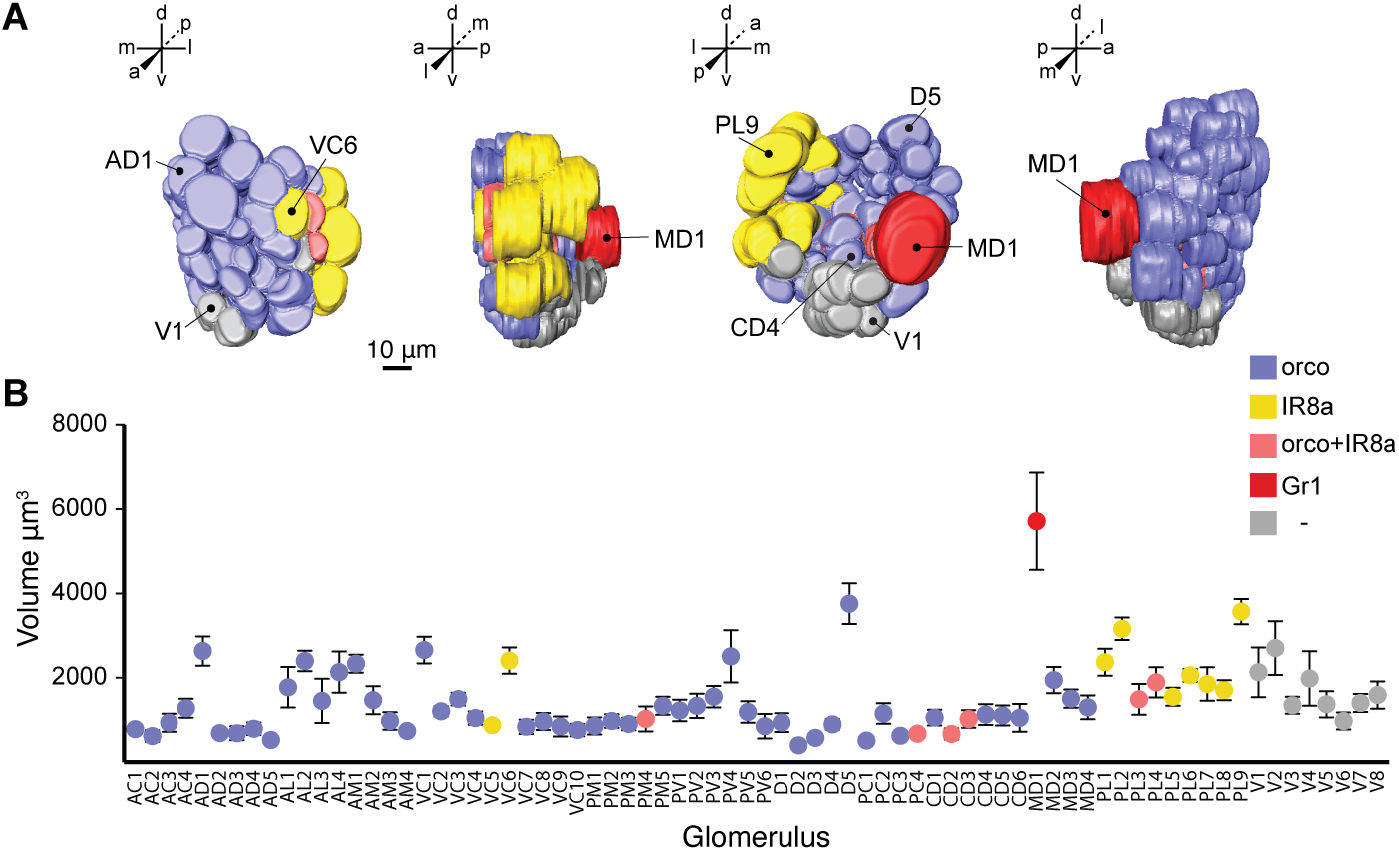
CO_2_ receptor complex neurons innervate the largest glomerulus in the *Aedes aegypti* antennal lobe. **(A)** 3D reconstructed model of the left antennal lobe of a female *A. aegypti* mosquito as seen from the anterior, lateral, posterior and medial perspectives. Template genotype for model: *orco^QF2^ > 30XQUAS-mCD8::GFP*. Landmark glomeruli are indicated. **(B)** Glomerular volumes from the female left antennal lobe. Mean volumes ± s.e.m. from spatially invariant glomeruli are plotted, *n* = 7 brains. Volumes varied significantly (one-way ANOVA, *P*< 0.0001). Tukey’s Multiple Comparison Test, *P* < 0.05 for all comparisons to the CO_2_ receptor glomerulus MD1 labeled in red.

Markedly increased glomerular subdivision of the *A. aegypti* antennal lobe relative to *Drosophila* (Shankar & McMeniman, 2020) makes assigning odor-evoked neurophysiological responses in the mosquito a daunting challenge. To overcome this, we leveraged the detailed receptor-to-glomerulus antennal lobe reference maps we generated, and applied functional imaging with calcium modulated photoactivatable ratiometric indicator (CaMPARI2) (Moeyaert et al., 2018) to map patterns of CO_2_ and L-(+)-lactic acid evoked activity in the antennal lobe. This genetically encoded, ratiometric calcium indicator photoconverts from green to red when simultaneously exposed to 405nm light and high levels of calcium (Fosque et al., 2015; Moeyaert et al., 2018), and has previously been applied for activity-dependent neural labeling only in model organisms such as mice, zebrafish and flies.

To pilot the efficacy of CaMPARI2 to map odor-evoked activity across glomeruli of the *A. aegypti* antennal lobe, we first calculated photoconversion ratios in the axon terminals of *Gr1* (+) OSNs innervating the MD1 glomerulus in response to stimulation with CO_2_ and synthetic air (*Figure 4A–B*) using *Gr1^QF2^ > 30XQUAS-CaMPARI2* female mosquitoes that we engineered. For CaMPARI2 photoconversion assays, live head-tethered mosquito preparations with surgically exposed antennal lobes were stimulated with simultaneous pulses of 1% CO_2_ and 405 nm light. Replicate brain samples were subsequently dissected and antennal lobe glomeruli rapidly co-stained with fluorophore-conjugated phalloidin to facilitate spatial mapping of CaMPARI2 signal using confocal microscopy of the antennal lobe (*Figure 4—figure supplement 1*). Encouragingly, MD1 exhibited a significantly higher rate of CaMPARI2 photoconversion in CO_2_-stimulated mosquitoes versus those that were stimulated with synthetic air (*Figure 4C*), validating the ability of 405nm light to penetrate mosquito brain tissue and photoconvert CaMPARI2 in an activity-dependent fashion in glomeruli such as MD1 positioned deep below the antennal lobe surface (Shankar & McMeniman, 2020).

**Figure 4.**
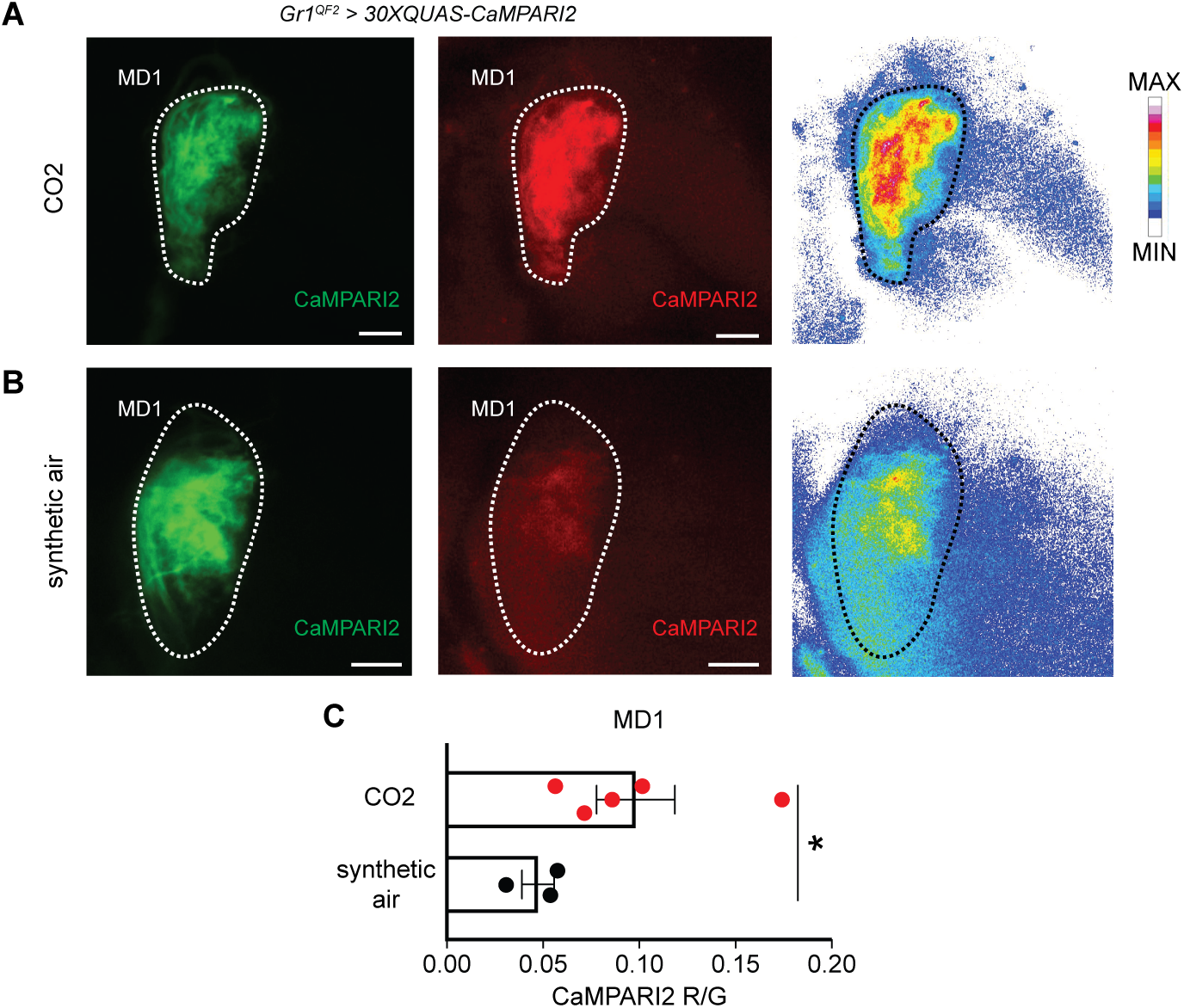
The *Aedes aegypti* MD1 glomerulus detects CO_2_. (**A** to **B**) CaMPARI2 green and red fluorescence in the *Gr1* (+) MD1 glomerulus after stimulation with (A) CO_2_, and (B) synthetic air. Right panels are heatmaps of red fluorescence intensity. MD1 from the left antennal lobe was imaged at 63X magnification. Scale bars: 10 μm. **(C)** CaMPARI2 photoconversion values in MD1, mean R/G values ± s.e.m. plotted, Mann-Whitney test *P* = 0.037 *.

We next tested whether synergy between the sweat odorant L-(+)-lactic acid and CO_2_ could be detected in specific *IR8a* (+) glomeruli of *IR8a^QF2^ > 30XQUAS-CaMPARI2* female mosquitoes, given that IR8a (Raji et al., 2019) and the CO_2_ receptor pathways (McMeniman et al., 2014) are both required for synergistic attraction of *A. aegypti* to these two odorants. Strikingly, when CO_2_ was coapplied with L-(+)-lactic acid to mosquitoes we observed strong CaMPARI2 photoconversion in specific *IR8a* (+) glomeruli (*Figure 5A*). In contrast, application of L-(+)-lactic acid, CO_2_ or synthetic air alone only yielded weak photoconversion across *IR8a* (+) glomeruli (*Figure 5B–D*). In particular, two out of the twelve *IR8a* (+) glomeruli we mapped, PL5 and PL6, exhibited significant differences in mean CaMPARI2 photoconversion ratios when co-stimulated with CO_2_ and L-(+)-lactic acid, compared to L-(+)-lactic acid or CO_2_ alone and synthetic air controls (*Figure 5E–F; Figure 5—figure supplement 1* and *Figure 6A–D*).

**Figure 5.**
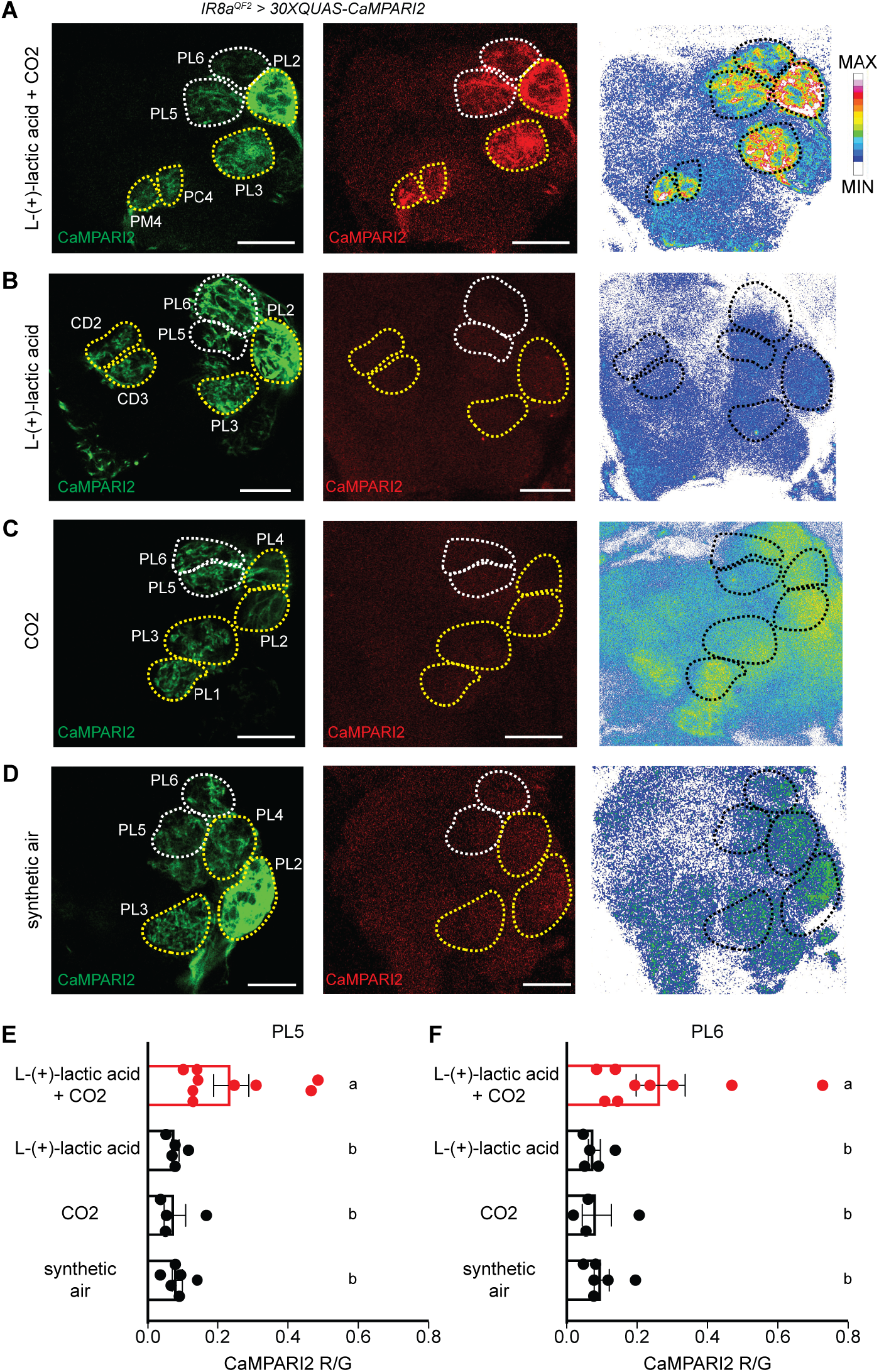
CO_2_ and L-(+)-lactic acid co-stimulation synergistically gates pre-synaptic calcium levels in acid-sensing *IR8a* neurons. (**A** to **D**) CaMPARI2 green and red fluorescence in *IR8a* (+) glomeruli after stimulation with (A) L-(+)-lactic acid + CO_2_, (B) L-(+)-lactic acid, (C) CO_2_ and (D) synthetic air. Right panels are heatmaps of red fluorescence intensity. Dotted outlines represent the boundaries of all *IR8a* (+) glomeruli in these representative z-slices. Left antennal lobe imaged at 63X magnification. Scale bars: 10 μm. (**E** to **F**) CaMPARI2 photoconversion values in (E) PL5 and (F) PL6 glomeruli, mean R/G values ± s.e.m. plotted, Tukey’s Multiple Comparison Test, *P* < 0.05 for all comparisons to ‘L-(+)-lactic acid + CO_2_’ in both glomeruli.

**Figure 6.**
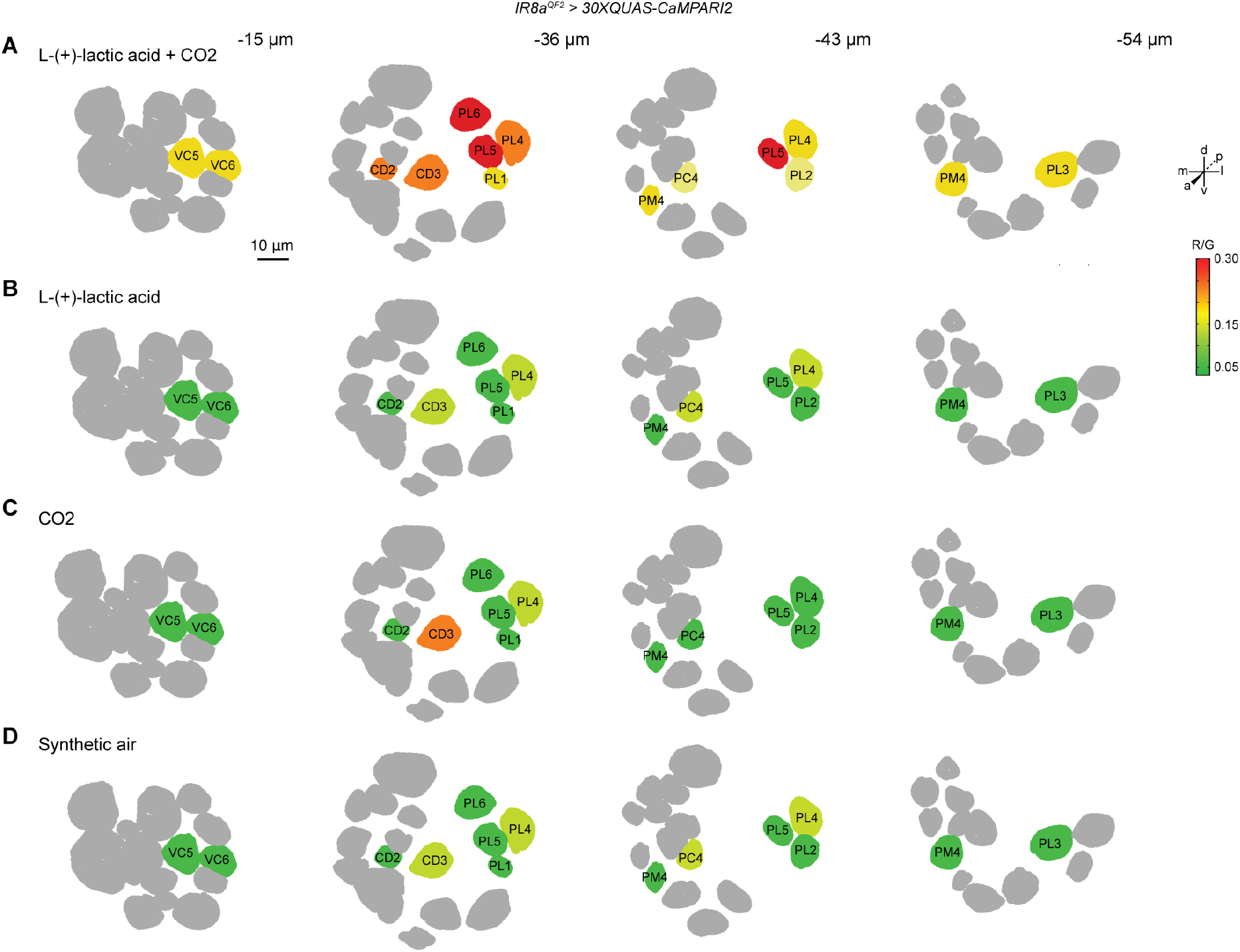
CaMPARI2 mapping illuminates synergistic coding of CO_2_ and L-(+)-lactic acid in *IR8a* (+) glomeruli PL5 and PL6 of the *Aedes aegypti* antennal lobe. Average CaMPARI2 photoconversion (R/G) values in *IR8a* (+) glomeruli from *IR8a^QF2^ > 30XQUAS-CaMPARI2* females in response to co-stimulatation with **(A)** L(+)-lactic acid + CO_2_, **(B)** L(+)-lactic acid alone, **(C)** CO_2_ alone and **(D)** synthetic air alone are shown. Photoconversion values for *IR8a* (+) glomeruli are plotted as a heat map on the phalloidin reference atlas. Four antennal lobe slices highlighting the complement of *IR8a* (+) glomeruli screened are shown, with each slice increasing in depth from the left to right of each panel. Scale bar: 10 μm.

## Discussion

The human sweat odorant L-(+)-lactic acid alone is unattractive to female *A. aegypti* (Acree et al., 1968). However, this molecule powerfully synergizes with the breath volatile CO_2_ to elicit mosquito behavioral attraction when these two olfactory stimuli are combined together (Acree et al., 1968; Eiras & Jepson, 1991; McMeniman et al., 2014; Smith et al., 1970). Using neural activity mapping with the ratiometric fluorescent sensor CaMPARI2 (Moeyaert et al., 2018), here we provide evidence for a circuit integrating CO_2_ and L-(+)-lactic acid detection that operates at the level of the first olfactory synapse in the mosquito antennal lobe.

To map glomeruli in the *A. aegypti* antennal lobe responsive to L-(+)-lactic acid and CO_2_, we first sought to gain genetic access to OSNs detecting these human-related odorants. To do this, we used CRISPR-Cas9-mediated homologous recombination to insert *T2A-QF2* in-frame fusion cassettes (Diao & White, 2012; Matthews et al., 2019) into the chemoreceptor co-receptor loci *IR8a* and *Gr1*. These two genes are expressed separately in OSNs that detect L-(+)-lactic acid and CO_2_, respectively (Jones et al., 2007; McMeniman et al., 2014; Raji et al., 2019). For neuroanatomical comparison, we also generated a *T2A-QF2* in-frame fusion in the *orco* gene (Larsson et al., 2004), given this co-receptor is highly expressed across *A. aegypti* olfactory tissues (Matthews et al., 2016). Using this genome engineering approach, we succeeded in developing a suite of OSN-specific *T2A-QF2* driver lines that facilitate transgenic labeling of the vast majority (~80%) of *A. aegypti* antennal lobe glomeruli.

During optimization of these transgenic reagents, we determined that *3xP3* markers typically used in insect transgenesis (Berghammer et al., 1999) induced both spurious reporter and background fluorescence in the *A. aegypti* brain central brain when integrated as part of the *T2A-QF2* in-frame fusion cassettes at these three target genomic loci. Of note, removing these *3xP3* markers by *Cre-LoxP* mediated cassette excision (Häcker et al., 2017) to generate marker-free *T2A-QF2* driver lines solved this problem. This strategy facilitated definitive assignment of receptor-to-glomerulus innervation patterns for each target chemoreceptor, when one copy of each marker-free *T2A-QF2* driver was complexed with two copies of a *15XQUAS-mCD8::GFP* responder. This approach strongly labeled OSN axonal processes and terminals demarcating each glomerulus in the antennal lobe.

Using these optimized transgenic tools and immunohistochemistry, we confirmed our previous observations using neural staining (Shankar & McMeniman, 2020) that the *A. aegypti* antennal lobe contains ~ 80 total glomeruli. Of these, we could reliably identify 74 spatially conserved glomeruli by cross-referencing nc82 and GFP channels to demarcate their boundaries. Across replicate brain samples, we consistently annotated the same complement of *IR8a* (+), *Gr1* (+) and *orco* (+) glomeruli in the antennal lobe. We determined that *IR8a* (+) OSNs innervate 15 glomeruli mostly localized to the posterolateral region of the *A. aegypti* antennal lobe, while *Gr1* (+) OSNs innervated a innervate a single, posteriorly positioned glomerulus in the mediodorsal antennal lobe region called MD1 (Shankar & McMeniman, 2020). Consistent with a previous study that employed a promoter-*QF2* fusion strategy to label *orco* (+) OSNs in *Anopheles gambiae* (Riabinina et al., 2016), we determined using a *T2A-QF2* in-frame fusion that *orco* (+) OSNs innervate a major proportion (at least 56) of all antennal lobe glomeruli in *A. aegypti*, as well as projecting to the taste center of the mosquito brain known as the subesophageal zone. This suggests that *orco* (+) neurons may have conserved roles in olfaction and taste in these two major mosquito disease vectors.

Using spatial mapping, we estimate that a small subset of six glomeruli in the *A. aegypti* antennal lobe co-express *IR8a* and *orco* based on overlapping patterns of glomerular labeling in this brain center. Indeed, immunostaining experiments with complementary *T2A-QF2* driver reagents developed by others (Younger et al., 2020) have revealed co-expression of *orco* and *IR8a* in a small number of OSNs from the *A. aegypti* antennae. Use of intersectional genetic reagents such as the split-*QF* system (Riabinina et al., 2019) and multiplexed whole mount in situ RNA hybridization for olfactory tissues recently developed for *A. aegypti* (Younger et al., 2020) will help to further refine the receptor-to-glomerulus maps described here for *orco*, *IR8a* and *Gr1*, towards mapping to the expansive array of other chemoreceptors expressed with these co-receptors in the *A. aegypti* olfactory system (Matthews et al., 2018; Matthews et al., 2016).

Reduced or more variable numbers of antennal lobe glomeruli and receptor-to-glomerulus labeling patterns using *T2A-QF2* in-frame fusions in *A. aegypti* have been concurrently described by two complementary studies (Younger et al., 2020; Zhao et al., 2020). We suggest our use of marker-free *T2A-QF2* driver strains and a cumulative 30 copies of membrane-tethered GFP to label OSNs, may have facilitated a clearer delineation of glomerular boundaries within the *A. aegypti* antennal lobe. Our optimized pipeline for generating *T2A-QF2* in-frame fusions and the reporter strains developed here may therefore be broadly useful for future studies of neuroanatomy and odor-coding in the *A. aegypti* antennal lobe. Application of these tools may also facilitate cell-type specific genetic access to other target genes in the mosquito nervous system and other tissues, without the need to identify enhancers and promoters to drive reporter expression.

Exposure to CO_2_ strongly activates mosquito flight (Eiras & Jepson, 1991) and enhances both the kinetics and fidelity of olfactory responses from *A. aegypti* towards whole human scent and several carboxylic acids, including L-(+)-lactic acid (Carlson et al., 1973; Dekker & Cardé, 2011; Dekker et al., 2005). Additionally, detection of CO_2_ is synergistically integrated with sensation of warmth and visual contrast to drive robust behavioral attraction to these combined stimuli in *A. aegypti* (Burgess, 1959; McMeniman et al., 2014; van Breugel et al., 2015; Vinauger et al., 2019; Zhan et al., 2021). Presumably such multimodal integration of CO_2_ and other sensory cues acts to enhance reliable localization of human hosts (Cardé, 2015). Leveraging the detailed receptor-to-glomerulus maps developed here, we confirmed that the CO_2_ receptor glomerulus MD1 is the largest cluster of neuropil in the *A. aegypti* antennal lobe (Shankar & McMeniman, 2020). The large volume of MD1 may indicate extensive synaptic connectivity between CO_2_ receptor neurons, local interneurons innervating other glomeruli in the antennal lobe, and projection neurons innervating higher order olfactory processing centers of the mosquito brain. This likely reflects the critical importance of CO_2_ to multiple facets of *A. aegypti* host-seeking behavior.

Our CAMPARI2 mapping data in the *A. aegypti* antennal lobe indicates that presynaptic calcium is significantly elevated in *IR8a* (+) OSN axon terminals innervating PL5 and PL6 glomeruli when mosquitoes are co-stimulated with CO_2_ and L-(+)-lactic acid. We propose such increases in pre-synaptic calcium concentration gate synaptic transmission from L-(+)-lactic acid sensitive OSNs to higher order neurons that orchestrate downstream behavioral attraction to these combined stimuli. Given the silent nature of *IR8a* (+) glomerular responses to either ligand alone observed with CaMPARI2, we hypothesize that this binary synergism may occur via disinhibitory local circuitry (Zhao et al., 2019) operating between the CO_2_-sensitive glomerulus MD1 and axon terminals of these lactic acid-sensitive *IR8a* (+) glomeruli in the antennal lobe.

At the synapse, presynaptic calcium levels may be modulated over varying timescales via neuromodulators (Higley & Sabatini, 2010; McKay et al., 2007). Intriguingly, transient optogenetic activation of *A. aegypti* CO_2_ neurons mimicking a fictive pulse of this gas, can yield a persistent behavioral state and attraction to warmth for more than 10 min (Sorrells et al., 2021), suggesting that CO_2_ detection may evoke neuromodulation. In addition to applying optogenetics, development of techniques for cell-type specific mutagenesis (Poe et al., 2019; Xue et al., 2014) in *A. aegypti* may further assist to query the role of candidate neuropeptides, neuropeptide receptors or other target genes in mediating synergism between CO_2_ and L-(+)-lactic acid detection in the antennal lobe. Circuit tracing (Ruta et al., 2010) may also assist to map neural connectivity between the CO_2_ receptor glomerulus MD1 and the *IR8a* (+) glomeruli sensitive to L-(+)-lactic acid described here, or other sensory processing centers of the mosquito brain.

In various organisms ranging from *C. elegans* to mice, a multitude peripheral and central mechanisms exist to modulate the salience of individual or blends of olfactory cues (Ko et al., 2015; Su et al., 2012; Tsunozaki et al., 2008; Wilson & Laurent, 2005; Xu et al., 2020; Yaksi & Wilson, 2010; Zak et al., 2020). Extracellular recordings from the dendrites of L-(+)-lactic acid sensitive neurons on the *A. aegypti* antennae (Davis & Sokolove, 1976), however, indicate that CO_2_ and L-(+)-lactic acid do not synergize together at the receptor or olfactory sensillum levels of olfactory coding.

Upon detection of CO_2_, *A. aegypti* must reliably track host volatiles including L-(+)-lactic acid emanating from a human host in order to obtain a blood meal. As byproducts of cellular respiration in different contexts, both CO_2_ and lactic acid are incredibly common molecules found in nature – challenging the mosquito with a noisy olfactory environment. In other insect species such as *Drosophila*, cross-talk between olfactory glomeruli can occur via the action of excitatory and inhibitory local interneurons, modulating the gain control of olfactory circuits (Chou et al., 2010; Hong & Wilson, 2015; Liu & Wilson, 2013; Nagel & Wilson, 2016; Wilson & Laurent, 2005; Yaksi & Wilson, 2010). Intra-glomerular communication between CO_2_ and L-(+)-lactic acid sensitive neurons in the *A. aegypti* antennal lobe may similarly serve to act as a coincidence detector and filter that allows mosquitoes to quickly process instructive host-related olfactory information during their quest for a human host.

Rapid feedback between antennal lobe glomeruli to yield disinhibition of OSN axon terminals may therefore represent a simple, yet flexible circuit for this prolific disease vector to faithfully identify signature combinations of human odorants and navigate their environment to improve the fidelity of their hunt for humans. Given that multiple human odorants in combination likely lie at the heart of mosquito lust for human scent (Bosch et al., 2000; Chen et al., 2019; Geier et al., 1999), further dissection of synergistic patterns of odor coding operational within the mosquito antennal lobe may reveal key human volatiles and chemosensory circuitry that can be targeted to combat mosquito-borne diseases such as dengue, Zika and malaria.

## Materials and Methods

### Mosquito Stock Maintenance

The *Aedes aegypti LVPib12* strain (Nene et al., 2007) was used as the genetic background for generation of all transgenic lines and subsequent assays. Mosquitoes were maintained with a 12 hr light:dark photoperiod at 27°C and 80% relative humidity using a standardized rearing protocol (Shankar & McMeniman, 2020). All experiments were conducted with mated, non-blood fed *A. aegypti* females that were 5-10 days old. Adult mosquitoes were provided constant access to a 10% w/v sucrose solution. Stock and composite genotypes used in each figure panel are detailed in Table 1.

**Table 1.**
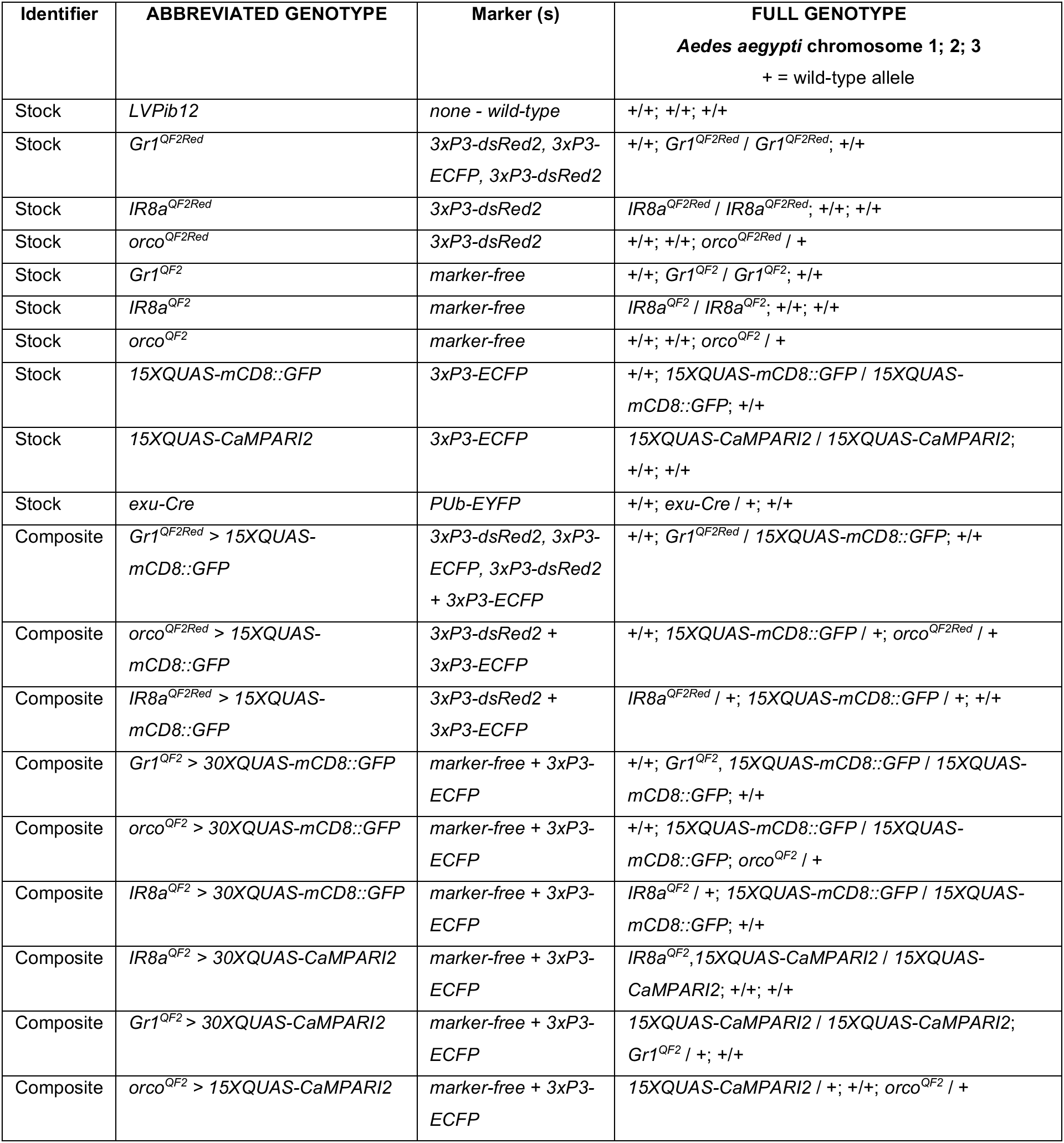
Complete genotypes of *Aedes aegypti* stocks and composite genotypes used in this study.

### Selection and *in vitro* transcription of sgRNAs

Single guide RNA (sgRNA) target sites in the coding sequences of *orco* (AAEL005776), *IR8a* (AAEL002922) and *Gr1* (AAEL002380) were identified using online design pipelines at http://zifit.partners.org/ZiFiT/ and http://crispr.mit.edu/. Candidate sgRNAs at each locus were prioritized for downstream use based on their putative lack of off-target activity in the *A. aegypti* genome. sgRNAs were transcribed and purified according to the method of Kistler et al. (2016) (Kistler et al., 2015). Briefly, DNA templates for sgRNA synthesis were generated by PCR with two partially overlapping PAGE-purified oligos (IDT) for each target. sgRNA was subsequently produced using the MegaScript T7 *in vitro* transcription kit (Ambion) and purified using the MEGAclear transcription clean-up kit (Invitrogen). Prior to microinjection, sgRNA activity was confirmed by *in vitro* cleavage assays with purified recombinant Cas9 protein (PNA Bio, Inc., CP01-200) following the manufacturer’s instructions. See Table S1 for final sgRNA sequences.

### *T2A-QF2* Donor Constructs

A base *T2A-QF2* donor construct (*pBB*) for CRISPR-Cas9 mediated homologous recombination into target chemoreceptor loci in *A. aegypti* was generated by sequential rounds of In-Fusion cloning (TakaraBio). This construct was generated in a pSL1180 backbone with an EcoNI site for in-frame fusion of a 5’ homology arm from a target gene with *T2A-QF2,* and BssHII site for insertion of the 3’ homology arm. To survey for homology arms, genomic DNA regions spanning each target site were first PCR amplified with CloneAmp (TakaraBio) using the following primers for *orco* (5’-TGCAAGTGGATCATTTGTCG-3’ and 5’-GTGCAATTGTGCCATTTTGA-3’), *IR8a* (5’-CAAAGTATAATTTCGCCCCCTCC-3’ and 5’-CTCTATGGCAGCCAAGATATTGG-3’) and *Gr1* (5’-AAGCCAGCTGGAAGGACATA-3’ and 5’-ACCGTTTGGAGGTTGAATTG-3’). PCR products were cloned into pCR2.1-TOPO (Invitrogen) for subsequent sequence verification. After determining the most common sequence clone for each region, homology arms flanking the CRISPR-Cas9 cut site were PCR-amplified and inserted into the *pBB* donor at the EcoNI site (5’ arm) and BssHII site (3’ arm) using the In-Fusion primers listed in Table S2, to generate a T2A in-frame fusion into the coding exon of interest. Three donor constructs that yielded successful integrations at these target loci included *pBB-AaOrco, pBB-AaIR8a* and *pBB-AaGr1*. Each *T2A-QF2* donor construct included a floxed *3xP3-DsRed2* transformation marker, as well as a *3xP3-ECFP* marker in the vector backbone outside the HDR cassette. This latter *3xP3-ECFP* marker was used to assess putative ends-in recombination events at the target locus or alternate off-target integrations elsewhere in the genome. *orco^QF2Red^* and *IR8a^QF2Red^* cassettes inserted in-frame as expected via ends-out recombination events. The *Gr1^QF2Red^* cassette inserted in-frame, but incorporated a duplicated copy of the plasmid backbone downstream of the *T2A-QF2* in-frame fusion via an ends-in recombination event.

### *Mos1 mariner QUAS* Reporter and Germline *Cre* Constructs

*QUAS* reporter and germline *Cre* cassettes were generated by sequential rounds of In-Fusion cloning (TakaraBio) into template plasmid backbones for *Mos1 mariner* transposition (Coates et al., 1998) as outlined in Table S3. All *QUAS* reporter constructs included a *3xP3-ECFP* transformation marker. The *pMOS* backbone for *15XQUAS-CaMPARI2* was modified to include a floxed *3xP3-ECFP* marker, and the *pMOS* backbone for *exu-Cre* was modified to have a *Polyubiquitin-EYFP* marker using standard cloning methods. Final plasmids that yielded transformants included: *pMosECFP-15XQUAS-mCD8::GFP, pMos-loxP-ECFP-loxP-15XQUAS-CaMPARI2* and *pMosEYFP-exu-Cre*.

The complete nucleotide sequences for all donor plasmids, *pMOS* vector backbones and *Mos1* helper (Coates et al., 1998) plasmids used in this study will be deposited to Addgene. Template materials are listed in Table S3.

### Generation of Transgenic Lines

*T2A-QF2* knock-in lines into *orco, IR8a* and *Gr1* were generated via CRISPR-Cas9 mediated homologous recombination (Kistler et al., 2015) using embryonic microinjection.

To generate the *Gr1^QF2Red^* insertion, an injection mixture consisting of sgRNA (40ng/ul), purified recombinant Cas9 protein (PNA Bio, 300ng/ul) and donor plasmid (500ng/ul) was prepared in microinjection buffer (5 mM KCl and 0.1 mM NaH_2_PO_4_, pH 7.2) and microinjected into the posterior pole of pre-blastoderm stage *LVPib12* embryos at the Insect Transformation Facility at University of Maryland (UM-ITF) using standard methods.

To generate the *orco^QF2Red^* and *IR8a^QF2Red^* insertions, sgRNA (100ng/ul) was mixed with the T2A-QF2 donor construct (100 ng/ul) for each target and microinjected into the posterior pole of transgenic *A. aegypti* pre-blastoderm stage embryos expressing *Cas9* under the maternal germline promoter *exuperantia* (Li et al., 2017). Transformed G_1_ larvae from all knock-in lines were isolated via the visible expression of *3xP3-DsRed2* marker in eye tissue. Transgenics were outcrossed to the *LVPib12* wild-type line for at least five generations prior to attempting to generate homozygous strains. Precise insertion of each donor construct was confirmed by PCR amplification and subsequent Sanger sequencing of regions covering the homology arms and flanking sequences on either side of the insertion.

*QUAS* reporter and *exu-Cre* strains were generated by co-injecting each *pMOS* donor construct (500 ng/ul) with a pKhsp82 helper plasmid (300 ng/ul) expressing the *Mos1* transposase (Coates et al., 1998) to foster quasi-random integration into the genome. Embryo microinjections were carried out by UM-ITF using standard techniques. For *QUAS* reporters, the G1 lines selected for stock establishment were those that had the strongest *3xP3-ECFP* expression levels in the eyes and ventral nerve cord, indicative of responder loci accessible for neuronal expression.

### *Cre-LoxP* Mediated Excision of *3xP3* Fluorescent Markers

Immunohistochemistry with *orco^QF2Red^>15XQUAS-mCD8::GFP, IR8a^QF2Red^> 15XQUAS-mCD8::GFP* and *Gr1^QF2Red^>15XQUAS-mCD8::GFP* adult female brains revealed spurious red and green fluorescence throughout the central brain. This was particularly noticeable in glia, including fixed brains not subjected to anti-GFP staining, suggesting potential interference in the expected *QF2/QUAS* transactivation pattern at these loci. As all of our *T2A-QF2* insertions included a downstream fluorescent marker cassette containing the *3xP3* synthetic promoter (Berghammer et al., 1999), which is a multimerized binding site for the paired-box transcription factor *Pax6* involved in glial and neuronal development (Quiring et al., 1994; Suzuki et al., 2016), we suspected such aberrant expression patterns may be due to promiscuous *3xP3* enhancer activity operating at these genomic loci.

To abrogate this effect, we developed a strategy to excise the floxed *3xP3* fluorescent marker cassettes from our *QF2^Red^* strains via crossing these genotypes to the germline *Cre* recombinase strain (*exu-Cre*) that we engineered. To remove floxed *3xP3* marker cassettes, we crossed males of each *QF2* driver line (*IR8a^QF2Red^, orco^QF2Red^, Gr1^QF2Red^*) to females of the *exu-Cre* line we generated. We then screened F_1_ progeny for loss of the *3xP3* fluorescent markers. In the case of the *Gr1^QF2Red^* line, the duplicated marker due to the ends-in insertion was incompletely removed in F_1_ progeny, so progeny still containing visible *DsRed2* or *ECFP* markers were mated to their *exu-Cre* (+) siblings to ensure complete excision of all markers. Precise excision was confirmed for all three driver lines by PCR and Sanger sequencing. Using this approach, we successfully generated marker-free driver strains (*orco^QF2^, IR8a^QF2^* and *Gr1^QF2^*) which were devoid of all *3xP3* fluorescent markers and any apparent background fluorescence in the central brain.

### *Mos1 mariner* Splinkerette PCR

*QUAS* and *exu-Cre* transgenes inserted via *Mos1* mariner transposition were mapped to chromosomal locations (AaegL5.0 genome assembly) using a modified Splinkerette PCR (Potter & Luo, 2010). Genomic DNA from single transgenic individuals was digested using the restriction enzymes BamHI-HF, BglII, and BstYI (New England BioLabs) in separate reactions. Digests were left overnight (~16 hrs). BstYI reactions were subsequently heat-inactivated at 80°C for 20 minutes according to the recommended protocol. BamHI reactions were purified using the QIAquick PCR Purification Kit (QIAgen) according to manufacturer instructions and eluted in 50 μl H_2_O after 4 minutes of incubation at 50°C.

Digests of genomic DNA were ligated to annealed Splinkerette (SPLNK) oligos as described (Potter & Luo, 2010). SPLNK oligonucleotides 5’-GATCCCACTAGTGTCGACACCAGTCTCTAATTTTTTTTTTCAAAAAAA-3’ and 5’-CGAAGAGTAACCGTTGCTAGGAGAGACCGTGGCTGAATGAGACTGGTGTCGACACT AGTGG-3’ were first annealed and ligated to digested genomic DNA. The first- and second-round PCR amplification steps were modified, using the standard SPLNK oligos and new primers designed to the inverted repeat regions of the *Mos1 mariner* transposon. PCR products were amplified using Phusion High-Fidelity DNA Polymerase (NEB).

First round Splinkerette PCR was carried out using the primers 5’-CGAAGAGTAACCGTTGCTAGGAGAGACC-3’ and 5’-TCAGAGAAAACGACCGGAAT-3’ for the right inverted repeat, and 5’-CGAAGAGTAACCGTTGCTAGGAGAGACC-3’ and 5’-CACCACTTTTGAAGCGTTGA-3’ for the left inverted repeat. The second round of Splinkerette PCR was carried out using the primers 5’-GTGGCTGAATGAGACTGGTGTCGAC-3’ and 5’-TCCGATTACCACCTATTCGC-3’ for the right inverted repeat, and 5’-GTGGCTGAATGAGACTGGTGTCGAC-3’ and 5’-ATACTGTCCGCGTTTGCTCT-3’ for the left inverted repeat. For *QUAS-CaMPARI2*, the extension time of the second-round PCR was lengthened to 4 minutes to amplify longer segments of flanking DNA. PCR products were gel purified and Sanger sequenced with additional sequencing primers for the right (5’-AAAAATGGCTCGATGAATGG-3’) and left (5’-GGTGGTTCGACAGTCAAGGT-3’) inverted repeats. BLAST searches were used to map Splinkerette fragments derived from each *Mos1 mariner* cassette to coordinate locations in the genome at canonical TA dinucleotides (Richardson et al., 2009) and insertion sites (Table S4) were subsequently confirmed by PCR.

### Genotyping *Gr1^QF2Red^* and *Gr1^QF2^*

*Gr1^QF2Red^* and *Gr1^QF2^* knock-ins were genotyped using a multi-primer PCR assay with the forward primer: 5’-CATGTACATCCGCAAGTTGG-3’; and two standard reverse primers: 5’-TGTTAGTGAGATCAGCGAACCT-3’ and 5’-GATCAACCCACAGATGACGA-3’. Fragments for size-based genotyping were amplified via DreamTaq (Thermo Scientific) and analyzed by conventional agarose gel electrophoresis. Each of the reverse primers was used at half the normal concentration. This resulted in a single 689 bp amplicon in homozygous mosquitoes; a single 884 bp amplicon in wild-type mosquitoes; and two amplicons, one at 689 bp and one at 884 bp, in heterozygous mosquitoes.

### Genotyping *IR8a^QF2Red^* and *IR8a^QF2^*

*IR8a^QF2Red^* and *IR8a^QF2^* knock-ins were genotyped using a multi-primer PCR assay with the forward primer: 5’-AGGAGATTGCGCTTGTCCTA-3’; and two standard reverse primers: 5’-CCCCGACATAGTTGAGCATT-3’ and 5’-TGTTAGTGAGATCAGCGAACCT-3’. Each of the reverse primers was used at half the normal concentration. This resulted in a single 560 bp amplicon in homozygous mosquitoes; a single 501 bp amplicon in wild-type mosquitoes; and two amplicons, one at 560 bp and one at 501 bp, in heterozygous mosquitoes.

### Genotyping *orco^QF2Red^* and *orco^QF2^*

*orco^QF2Red^* and *orco^QF2^* knock-ins were genotyped using conventional PCR. The PCR reaction used the forward primer: 5’-GCGATAGCGTCAAAAACGTA-3’ and reverse primer: 5’-ATTCCTTGAAGGTCCATTGCAG-3’. This resulted in an 1842 bp amplicon corresponding to the *orco^QF2^* allele, a 3129 bp amplicon corresponding to the *orco^QF2Red^* allele, and a 367 bp amplicon corresponding to the wild-type allele. Heterozygotes had both wild-type and transgenic PCR bands.

### Genotyping *15XQUAS-mCD8::GFP*

*15XQUAS-mCD8::GFP* was genotyped using conventional PCR. The PCR reaction used the forward primer: 5’-TCCAGCCGATAGGAACAATC-3’ and reverse primer: 5’-CAAATCCGAATTTCCCGTAA-3’. This resulted in a single 5797 bp amplicon for homozygotes and a 444 bp amplicon for the wild-type allele. Heterozygotes typically only had the wild-type PCR band.

### Genotyping *15XQUAS-CaMPARI2*

*15XQUAS-CaMPARI2* was genotyped using a multi-primer PCR assay with the forward primer: 5’-GTTTGACCAAATGCCGTTTC-3’; and two standard reverse primers: 5’-GTCGATAGGCGCGTAGTGTA-3’ and 5’-CACCACTTTTGAAGCGTTGA-3’. Each of the reverse primers was used at half the normal concentration. This resulted in a single 645 bp amplicon in homozygous mosquitoes; a single 874 bp amplicon in wild-type mosquitoes; and two amplicons, one at 645 bp and one at 874 bp in heterozygous mosquitoes.

### Transgenic Stock Maintenance and Composite Genotypes

*Gr1^QF2Red^*, *Gr1^QF2^*, *IR8a^QF2Red^* and *IR8a^QF2^* driver lines were maintained as homozygous stocks. *orco^QF2Red^* was maintained as a heterozygous stock by outcrossing to *LVPib12* each generation. *orco^QF2^* was maintained as a heterozygous stock by outcrossing to either *LVPib12* or *QUAS-mCD8::GFP* each generation and screening for GFP fluorescence in olfactory tissues of the progeny. *15xQUAS-mCD8::GFP* and *15xQUAS-CaMPARI2* responder lines were maintained as homozygous stocks. The *exu-Cre* line was maintained as a heterozygous stock by outcrossing to *LVPib12* each generation.

### Immunohistochemistry

Immunostaining of female *A. aegypti* brains was performed as previously described (Shankar & McMeniman, 2020), with minor modifications. Briefly, severed mosquito heads were fixed in 4% paraformaldehyde (Milonig’s buffer, pH 7.2) for three hours after which brains were carefully dissociated from the head capsule, pigmented ommatidia and air sacs. Dissected brains were then subjected to three 20 min washes at room temperature in PBST (0.1M PBS with 0.25% Triton-X 100), and allowed to incubate overnight in a blocking solution consisting of 2% normal goat serum (NGS) and 4% Triton-X 100 in 0.1M PBS at 4°C. Brains were then washed three times for 20 min each in PBST and incubated for three days at 4°C in a primary antibody solution containing mouse anti-BRP (DSHB, nc82-s, AB_2314866, 1:50 v/v) targeting the pre-synaptic active zone protein Bruchpilot (Hofbauer et al., 2009) and rabbit anti-GFP (Invitrogen, A-6455, 1:100 v/v) targeting mCD8::GFP. Brains were then washed three times for 20 min each in PBST and incubated for 3 days at 4°C in a secondary antibody solution consisting of goat anti-mouse Cy3 (Jackson ImmunoResearch, AB_2338680, 1:200 v/v) and goat anti-rabbit Alexa Fluor 488 (Invitrogen, A-11008, 1:200 v/v). All primary and secondary antibody dilutions were prepared in PBST with 2% v/v NGS. Brains were finally washed three times for 20 min each in PBST at room temperature and mounted in 20 ul of Slow-Fade Gold antifade mountant (Invitrogen, S36936) on glass slides with coverslip bridges (Number 2-170 μm).

### Immunohistochemistry Image Acquisition Settings

Brain immunostaining images were acquired on a single-point laser scanning Carl-Zeiss LSM 780 confocal microscope. To capture images of the entire adult brain, a 10X objective lens (0.3 NA, Plan-Apochromat) was used. Excitation of Cy3 signal was achieved with a 561 nm solid-state laser line at 0.05 % laser power, and GaAsP detector gain set to 825. A 488 nm laser line was used to excite Alexa Fluor 488 (20% laser power, detector gain at 825). We additionally acquired images with a 20X objective lens (0.8 NA, Plan-Apochromat) to perform 3D reconstructions of the antennal lobes. For these the power of the 488 nm laser line was adjusted to 5%. For each antennal lobe, 60 z-slices with a z-step size of 1 μm and a 1024 × 1024-pixel resolution were acquired.

### Antennal lobe reconstructions

3D morphological reconstructions of left antennal lobes were performed as previously described (Shankar & McMeniman, 2020). Briefly, confocal images were imported into Amira (FEI Houston Inc) and then segmented by highlighting all pixels across a z-stack occupied by individual glomeruli. The nc82 (Bruchpilot) channel was used for manual segmentation of individual glomeruli. The GFP channel was then used to identify *orco, IR8a* and *Gr1* (+) glomeruli. Cross referencing signals obtained from nc82 and GFP channels within and between samples in this dataset helped to clearly delineate glomerular boundaries. 3D and 2D antennal lobe models were generated by surface rendering. The number of GFP labeled glomeruli in three replicate left antennal lobe reconstructions per genotype from *orco^QF2^ > 30XQUAS-mCD8::GFP, IR8a^QF2^ > 30XQUAS-mCD8::GFP* and *Gr1^QF2^ > 30XQUAS-mCD8::GFP* females were counted. The total number of glomeruli per lobe were counted in 7 of these samples: *orco^QF2^ > 30XQUAS-mCD8::GFP* (n=3)*, IR8a^QF2^ > 30XQUAS-mCD8::GFP* (n=3) and *Gr1^QF2^ > 30XQUAS-mCD8::GFP* (n=1).

### Glomerular volume and frequency

Glomerular volumes were obtained from the left antennal lobe using the nc82 channel. To name glomeruli, we first identified landmark glomeruli in each antennal lobe sample using a systematic *A. aegypti* antennal lobe reference key (Shankar & McMeniman, 2020). Each antennal lobe glomerulus labeled with GFP was named based on its spatial position relative to these landmarks and flanking glomeruli. We classified glomeruli as spatially ‘invariant’ or ‘variant’ based on their frequency of identification. A threshold frequency of 80% or more across reconstructions was designated for the classification of spatially invariant glomeruli. Glomerular volume and frequency values were calculated from a pooled dataset consisting of the 7 reconstructions where we named all constitutive glomeruli in the same left antennal lobe samples used for glomerular counts: *orco^QF2^ > 30XQUAS-mCD8::GFP* (n=3 brains)*, IR8a^QF2^ > 30XQUAS-mCD8::GFP* (n=3 brains) *and Gr1^QF2^ > 30XQUAS-mCD8::GFP* (n=1 brain).

For *Gr1^QF2^ > 30XQUAS-mCD8::GFP* reconstructions, we only fully named and obtained volume measurements for all constitutive glomeruli from one antennal lobe sample of this genotype. This was due to the fact a single, very large landmark glomerulus MD1 was GFP labeled across all *Gr1^QF2^ > 30XQUAS-mCD8::GFP* reconstructions. To validate assignment of this glomerulus as MD1, we annotated four flanking mediodorsal (MD) group glomeruli using the reference key in two additional replicate *Gr1^QF2^ > 30XQUAS-mCD8::GFP* reconstructions, and only observed GFP labeling in the glomerulus annotated as MD1.

To test whether mean volumes of invariant glomeruli differed significantly within this dataset, we performed a One-Way ANOVA and Tukey’s HSD post hoc test to correct for multiple comparisons.

### Imaging of Peripheral Olfactory Appendages

Live antenna, palp and proboscis tissue were dissected in 0.1M PBS and immediately mounted in Slow-Fade Gold antifade mountant (Invitrogen, S36936). Images were acquired on a Carl-Zeiss LSM 780 confocal microscope within 1 hour of dissection. To excite the *GFP* signal, the 488 nm laser line was used at 5% laser power. An additional DIC channel was used to visualize gross morphology of the peripheral tissue. Images of the antennae were acquired with a 20X objective lens (0.8 NA, Plan-Apochromat), while images of the palp and labella of the proboscis were taken with a 40X (1.3 NA, Plan-Apochromat) oil immersion objective.

### Live Mosquito Preparation for CaMPARI2 Photoconversion

To prepare mosquitoes for CaMPARI2 photoconversion (Moeyaert et al., 2018), mosquitoes were cold anesthetized and tethered to an imaging chamber. To do this, the thorax of a female mosquito was first affixed to the ventral surface of a 35mm petri dish lid (Eppendorf, 0030700112) using UV-curing adhesive (Bondic) immediately next to a 15mm diameter circular hole made in the lid center. Two additional drops of adhesive were applied to the ommatidia on the extremities of the mosquito head to prevent head movement. A small piece of clear tape (Duck EZ Start, Heavy Duty Packaging Tape) was then adhered over the center hole. The dorsal surface of the mosquito head was then gently affixed to the ventral adhesive tape surface covering the hole. An excised section of plastic coverslip (5mm x 3mm) was then affixed to the tape and used to shield the antennae from the adhesive tape surface and suspend these sensory appendages in the air.

The imaging chamber with head-fixed mosquito was then inverted and a rectangular incision approximately 400 μm X 200 μm was cut through the tape window where the dorsal head cuticle and ommatidia were affixed. The wide boundary of the incision was typically made immediately adjacent to the first antennal subsegment along the lateral-medial brain axis, while the short boundary of the incision extended along the dorsal-ventral brain axis. To create this window, segments of ommatidia and bridge cuticle between the left and right eyes were gently cut and excised using a surgical stab knife (Surgical Specialties Corporation, Sharpoint, Part # 1038016) to reveal the underlying antennal lobes. The exposed antennal lobes were then immediately immersed in an *A. aegypti* Ringer’s solution (Beyenbach & Masia, 2002) composed of 150 mM NaCl, 3.4 mM KCl, 5mM glucose, 1.8 mM NaHCO_3_, 1 mM MgCl_2_, 25mM HEPES and 1.7 mM CaCl_2_; pH 7.1. Mosquitoes were allowed to recover for a period of 15 min from cold anesthesia and surgery in a humidified chamber at room temperature prior to imaging.

### CaMPARI2 Photoconversion

For CaMPARI2 photoconversion, the tethered preparation was placed under a 20X water dipping objective (Olympus XLUMPLFLN20XW, 1.0 NA) ensuring that the antennal lobes expressing basal green CaMPARI2 signal were in focus. Each preparation was exposed to a combined photoconversion-odor stimulation regime consisting of repetitive duty cycles of four 500 ms pulses of 405nm light from an LED driver (Thorlabs, DC4104, 1000mA current setting) synchronized with a 1 s odorant pulse as previously outlined (Fosque et al., 2015), for 75 cycles with a total protocol duration of approximately 41 minutes.

### Odorant Delivery

Pulses of odorants were delivered using a custom olfactometer device (Lundström et al., 2010). 3mL of control (dH2O) or treatment (L-(+)-lactic acid solution, Sigma Aldrich, 27714) odorant solutions were placed into dedicated and sealed odor delivery vials. During ‘odor onset’, synthetic air (20% O_2_ - 80% N_2_, Airgas) at a flow rate of 1mL/s was passed through these delivery vials. Vial headspace was then piped via Teflon tubing into a carrier airstream of humified synthetic air that was directed at the olfactory appendages of the mosquito using a plastic pipette. During CaMPARI2 photoconversion assays, the tethered mosquito preparation always received a constant amount of airflow (5ml/s) during odor onset/offset from the stimulus pipette. This was achieved via solenoid valves simultaneously switching or combining humidified synthetic air, 5% CO_2_ (Airgas) and/or L-(+)-lactic acid headspace as required for different odor treatments. For trials involving CO_2_, a 1ml/s stream of 5% CO_2_ was diluted 1:5 into the carrier airstream for a final concentration at the specimen of 1% CO_2_.

### CaMPARI2 Sample Processing

Following photoconversion, the mosquito was gently untethered from the imaging chamber and the head severed and fixed in Milonig’s buffer for 20 minutes. The brain was then dissected out in calcium-free Ringer’s solution composed of 150 mM NaCl, 3.4 mM KCl, 5 mM glucose, 1.8 mM NaHCO_3_, 1 mM MgCl_2_, 25 mM HEPES and 10 mM EGTA. To stain glomerular boundaries, we incubated each brain in Alexa Fluor 647 Phalloidin (Invitrogen, A22287) prepared in calcium-free Ringer’s solution (1:40 v/v dilution) for 30 min. To prepare Alexa Fluor 647 phalloidin for use in imaging, first, a 400X DMSO stock solution was prepared according to the manufacturer’s instructions by dissolving the fluorophore in 150 ul of DMSO. 1 ul of this DMSO stock was diluted in 399 ul calcium-free Ringer’s solution to yield a 1X stock. This 1X stock was then further diluted 1:40 in calcium-free Ringer’s solution for staining. Brains were transferred directly from this solution into 20 ul of Slow-Fade Gold Antifade Mountant (Invitrogen, S36936) on glass slides with coverslip bridges (Number 2-170 μm) for CaMPARI2 and phalloidin imaging.

### CaMPARI2 Image Acquisition Settings

Antennal lobes from CaMPARI2 photoconversion assays were imaged with a 63X (1.4 NA) oil-immersion objective on a Zeiss 880, Airyscan FAST super-resolution single point scanning microscope. Excitation of red CaMPARI2 signal was achieved with a 561 nm solid-state laser line at 14 % laser power. Green CaMPARI2 was excited with a 488 nm argon laser line at 10% laser power. To visualize glomerular boundaries, a 633 nm diode laser was used to excite the Alexa-647 phalloidin fluorophore at 40% laser power. Master detector gain was set to a value of 800. We captured 0.987 μm z-slices of 1572 × 1572-pixel resolution in the FAST mode. Raw images were further processed by applying the Airyscan method with ‘auto’ processing strength.

### CaMPARI2 Image Analysis

Image analysis was carried out in Fiji (http://imagej.net/Fiji). We first applied a median filter (radius = 2 pixels) to remove noise and then a rolling ball subtraction (rolling ball radius = 80 pixels), to correct for non-uniformity of background intensities. We analyzed CaMPARI2 photoconversion in the left antennal lobe of all samples due the well-defined spatial arrangement and conspicuous boundaries of *IR8a* (+) glomeruli in this lobe with phalloidin straining. ROIs were defined by manually segmenting glomeruli using the free hand selection tool. We analyzed photoconversion ratios for 12/15 *IR8a* (+) glomeruli expressing cytosolic CaMPARI2 that could be reliably identified across all antennal lobe samples. These included: VC5, VC6, PL1-PL6, PM4, PC4, CD2 and CD3. The three remaining *IR8a* (+) glomeruli (PL7-PL9) are normally only labeled sparsely when expressing membrane-tethered mCD8::GFP, and cytosolic CaMPARI2 was not evident in them. The integrated density (mean gray value × area) for all z-slices of the ROI, which included all representative slices of a target glomerulus, was calculated in the green (488 nm) and red (560 nm) imaging channels. The final measure of photoconversion, the red to green ratio (R/G), was calculated as:

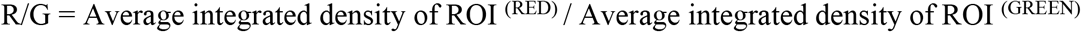

Glomeruli were named by co-localizing green and red CaMPARI2 signal to individual glomeruli evident in the Alex-Fluor 647 phalloidin channel and defining their spatial orientation relative to landmark and flanking glomeruli.

## Acknowledgments

We thank N. Kizito, B. Natarajan, W. Okoth, H. Rosado, G. Nasir, M. Gebhardt, V. Balta, A. Ellison and B. Burgunder for expert technical assistance; R. Harrell (UM-ITF) for mosquito embryonic microinjection services; S. Seo and A. Hammond for help with transgene mapping, C. Potter and E. Schreiter for constructs and technical advice; C. Huang, M. Schnitzer, B. Ferris, G. Maimon, C. Dan and V. Jayaraman for guidance on surgical preparations; and M. Wohl for comments on the manuscript. The mosquito template in Figure 1 was created for us by Biorender.com.

## Funding

This research was supported by funding from the National Institutes of Health NIAID (R21 AI139358-01), USAID (AID-OAA-F-16-00061) and Centers for Disease Control and Prevention (200-2017-93143) to C.J.M; and funding to G.M.T. as a postdoctoral fellow on The Molecular and Cellular Basis of Infectious Diseases (MCBID) Program (T32A1007417) from the NIH. O.S.A and M.L. were supported in part by a DARPA Safe Genes Program Grant (HR0011-17-2-0047), a DARPA ReVector program grant (HR0011-20-2-0030), and an NIH NIAID RO1 (RO1AI148300) awarded to O.S.A. The views, opinions and/or findings expressed should not be interpreted as representing the official views or policies of the Department of Defense or the U.S. Government. Microscopy infrastructure at Johns Hopkins School of Medicine Microscope Core Facility used in this research was supported by the National Institutes of Health NCRR (S10OD016374 and S10OD023548). We thank Terry Shelley at the JHU Center for Neuroscience Research Machine Shop for fabrication services supported by NINDS Center grant (NS050274). We further acknowledge generous support to C.J.M. from Johns Hopkins Malaria Research Institute (JHMRI) and Bloomberg Philanthropies. S.S. and G.M.T. were supported by JHMRI Postdoctoral Fellowships.

## Author Contributions

S.S. and C.J.M conceived the experimental design. M.L. and O.S.A. generated and provided the *exu-Cas9* strain. G.M.T., S.S. and C.J.M. assembled constructs for transgenesis and the custom olfactometer for odorant delivery. C.J.M. and G.M.T. screened, genotyped and maintained transgenic lines. E.D.S. performed confocal analyses of *IR8a* peripheral expression patterns. S.S. performed all other microscopy, immunohistochemistry, antennal lobe reconstructions, glomerular mapping and *CaMPARI2* imaging experiments. D.G. and S.S. analyzed the data. S.S. and C.J.M. drafted the manuscript.

## Competing interests

The authors declare no competing interests.

---

**Figure 1—figure supplement 1.**
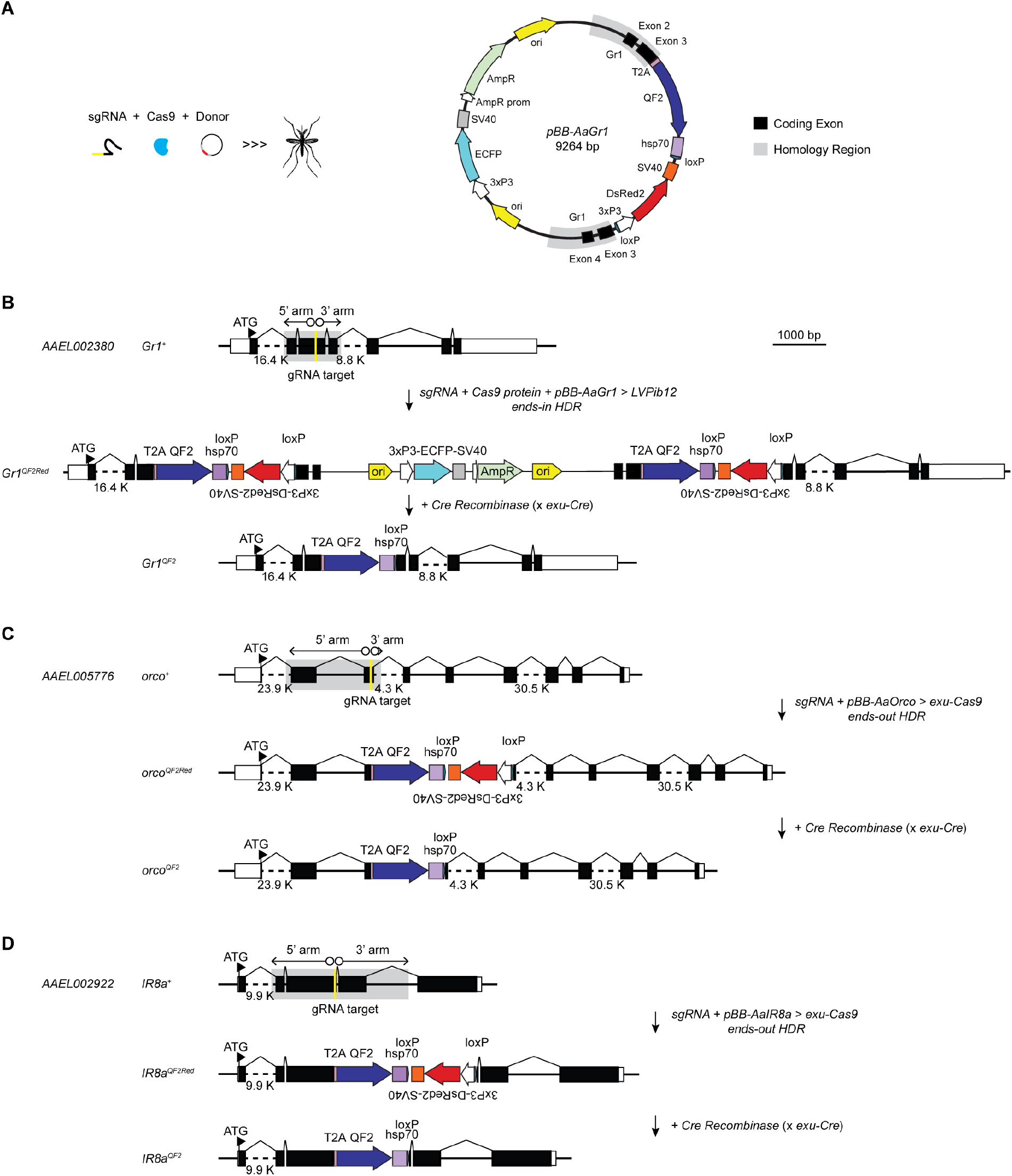
CRISPR-Cas9 mediated *T2A-QF2* in-frame fusion strategy to gain genetic access to broad OSN subsets in *Aedes aegypti*. **(A)** Mixtures of sgRNA, Cas9 protein and donor template (left) were injected in different formats into *A. aegypti* pre-blastoderm stage embryos to generate *T2A-QF2* in-frame fusion driver lines for *Gr1, orco* and *IR8a* genes. Lines were initially generated in *LVPib12* or *exu-Cas9 A. aegypti* genetic backgrounds. An example targeting construct for *Gr1* (right) is shown. **(B)** Schematic of the ends-in homology-directed repair (HDR) event that generated the *Gr1^QF2Red^* driver line. **(C-D)** Schematics of the ends-out HDR events that generated the *orco^QF2Red^* and *IR^QF2Red^* driver lines. The marker-free driver lines *Gr1^QF2^, Orco^QF2^* and *IR8a^QF2^* were generated by excising the floxed *3xP3-dsRed2* marker from these original integrations by crossing with an *exu-Cre A. aegypti* strain that we engineered and selecting for loss of the marker gene.

**Figure 1—figure supplement 2.**
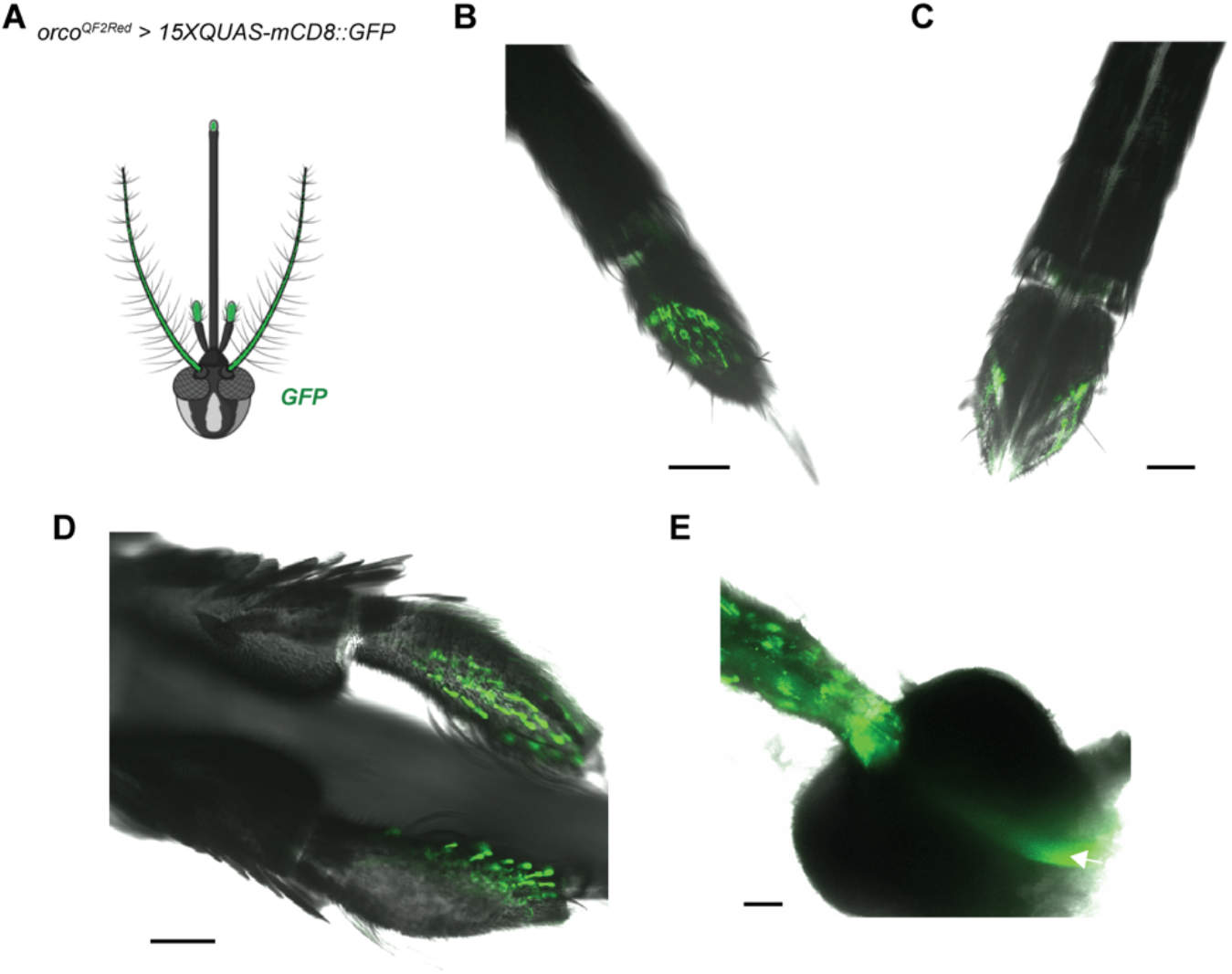
Innervation patterns of *orco* (+) olfactory sensory neurons across *Aedes aegypti* olfactory appendages. **(A)** Schematic of the mosquito head showing the gross expression pattern evident from *orco^QF2Red^ > 15XQUAS-mCD8::GFP* females in peripheral olfactory tissues. GFP labeling was observed in the antennae, maxillary palps and proboscis. **(B** to **C)** GFP expression from dorsal and ventral perspectives of the labella of the proboscis. Axonal projections of these neurons extend into the shaft of the proboscis. **(D)** *orco* expression was also strongly evident within capitate peg sensilla on the ventral surface of the maxillary palp. **(E)** Maximum intensity projection of a female pedicel. No *orco* expression was noted in the scolopidia of the pedicel while *orco* (+) axons in the antennal nerve (arrow) were observed to transect the pedicel. Scale bars: 50 μm.

**Figure 1—figure supplement 3.**
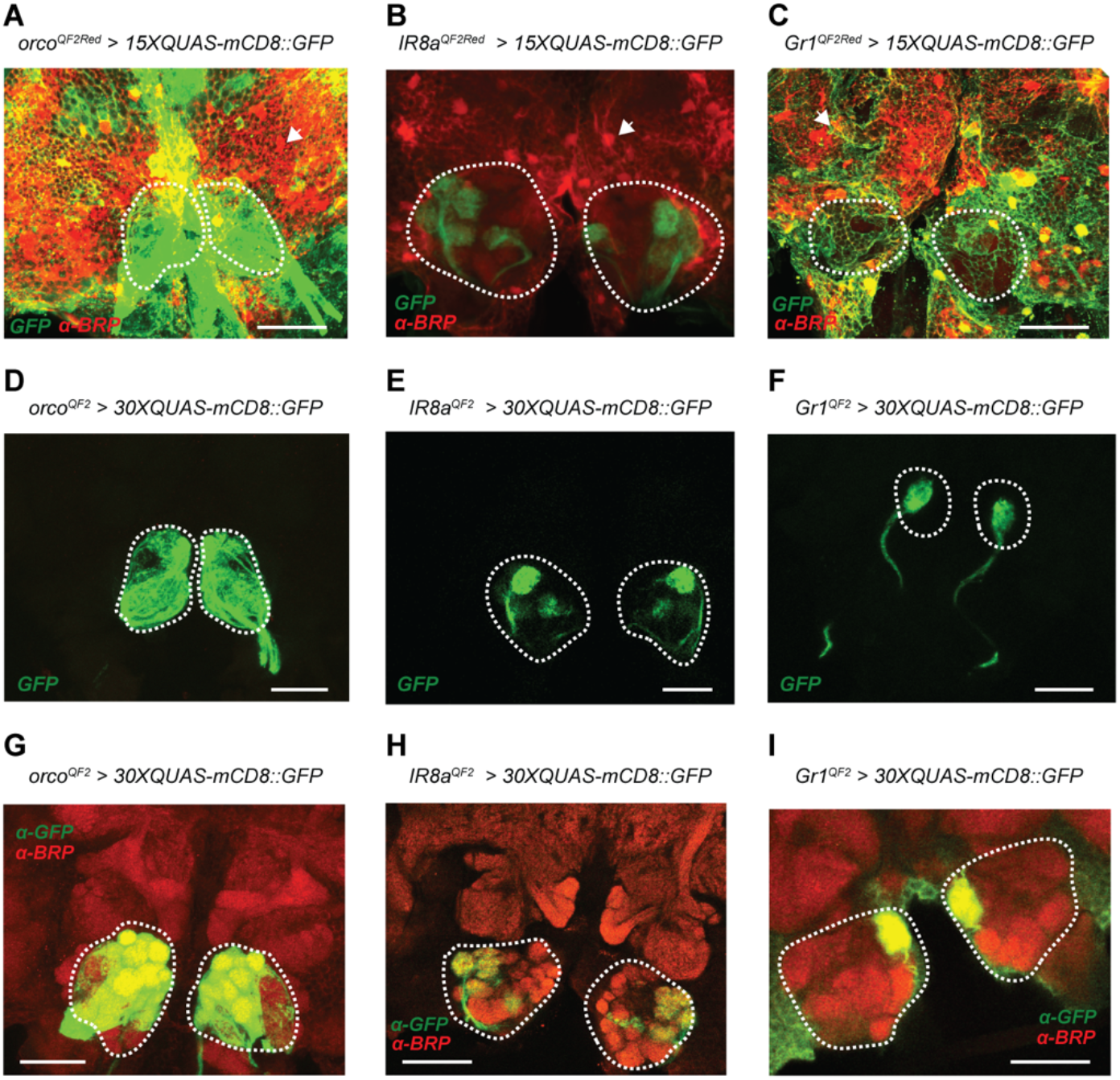
Promiscuous *QF2/QUAS* driven fluorescence in the *A. aegypti* brain and *Cre-LoxP* mediated excision of *3xP3* marker cassettes. The transgenic *QF2^Red^* driver lines described in this study carried a floxed *3xP3-DsRed2* cassette to mark successful transgenesis. However, during examination of the central brain we noted that this marker cassette was not only expressed in the optic lobes, but was also strongly expressed in hexagonal cells ensheathing the central brain which are putatively glia (arrows). In crosses involving the *orco^QF2Red^* and *Gr1^QF2Red^* driver lines, we also noted conspicuous green expression in these same locations suggesting the *3xP3* promoter was influencing the expression pattern of the integrated *T2A-QF2* transgene at these loci. **(A** to **C)** Anterior view of adult female mosquito brains at 20X magnification. Spurious *3xP3*-induced labeling occludes the antennal lobes and other neuropil, in each of the three driver lines, posing a significant challenge for confocal or two-photon imaging. **(D** to **F)** Genetic excision of this marker from the *QF2* driver lines alone markedly improved our ability to clearly view the antennal lobes in both unstained, and **(G** to **I)** immunostained brain preparations. Antennal lobe glomeruli are encircled by white dashed lines. Scale bars: 50 μm.

**Figure 2—figure supplement 1.**
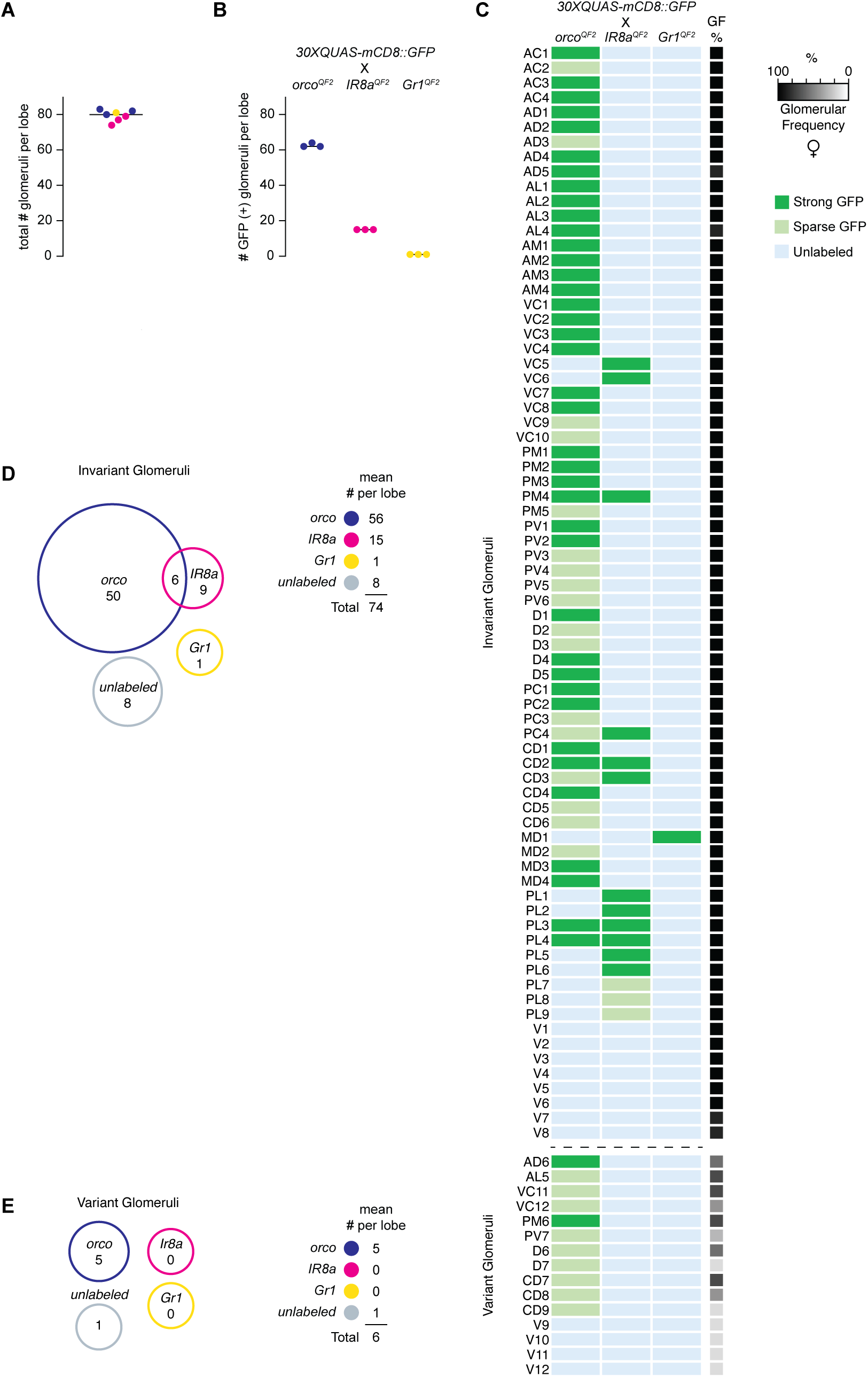
Classification and frequency of *Aedes aegypti* antennal lobe glomeruli. **(A)** Total number of glomeruli per reconstructed antennal lobe (n=7). **(B)** Number of GFP labeled glomeruli per genotype (n=3). **(C)** Innervation pattern of GFP-labeled *orco, IR8a* and *Gr1* (+) neurons in the *A. aegypti* antennal lobe. Glomeruli were either strongly or sparsely labeled with mCD8::GFP, or unlabeled. A heatmap of glomerular frequency (GF %) is shown in parallel. 74 spatially invariant glomeruli were characterized across reconstructions (frequency ≥ 80%). 15 variant glomeruli with different annotated names were also found across some reconstructions (frequency < 80%). **(D** to **E)** Venn diagrams summarizing the mean number of invariant and variant glomeruli, respectively that were GFP labeled by each chemoreceptor driver line.

**Figure 2—figure supplement 2.**
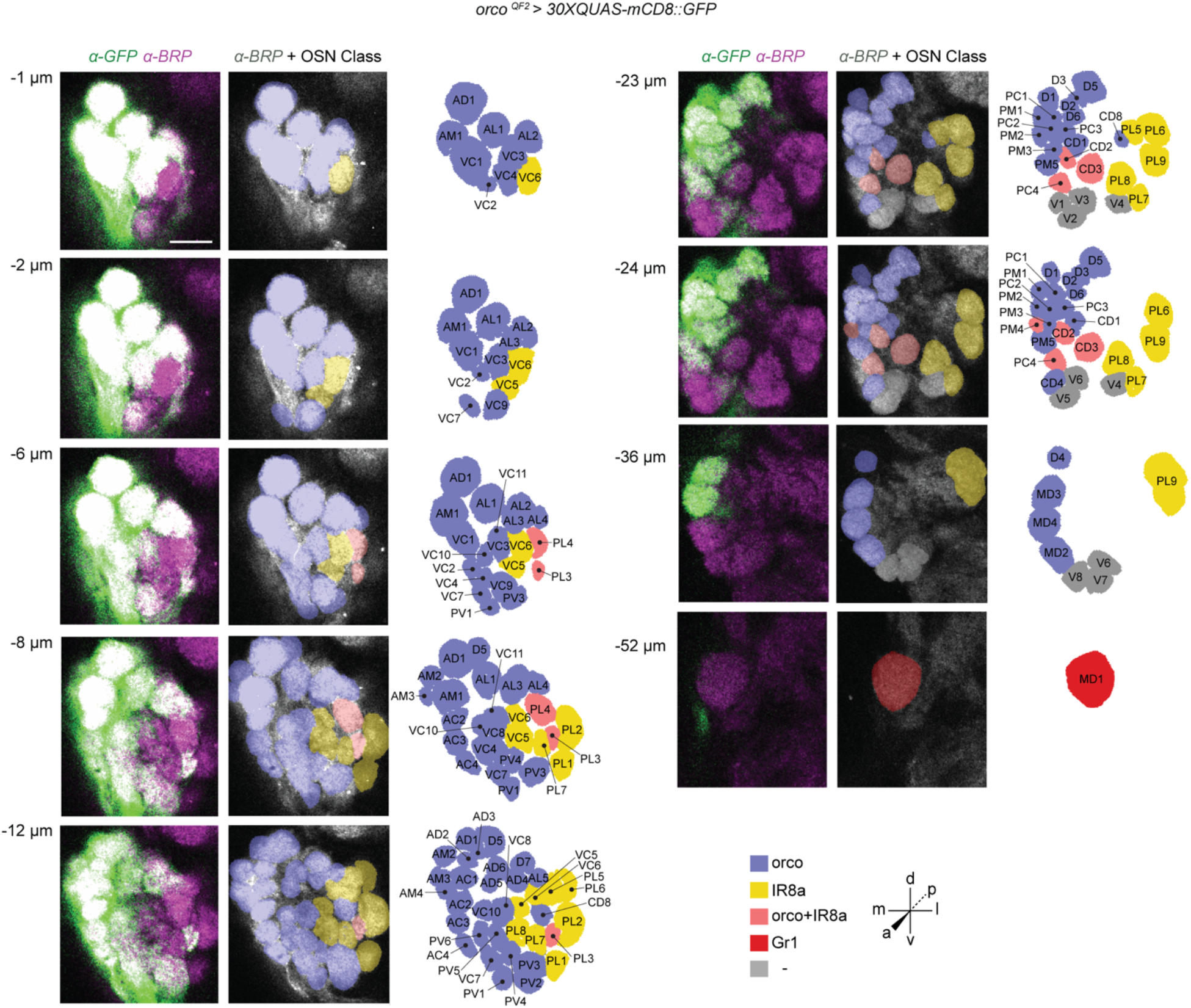
*orco* receptor to glomerulus 2D map. Representative confocal stack from the left antennal lobe of an adult female *orco^QF2^ > 30XQUAS-mCD8::GFP* mosquito. The left panels show the antennal lobe at 20X magnification, with the GFP signal from *orco* neurons in green and the neuropil stained with BRP (Bruchpilot) signal in magenta, to visualize glomerular boundaries. The middle panels show the BRP stained glomeruli overlayed with blue (*orco* +), yellow (*IR8a* +), pink (putatively *orco* + and *IR8a* +), red (*Gr1* +), and gray colors (unknown chemosensory identity). The right panels show detailed spatial maps of all antennal glomeruli in each slice. Nine representative focal planes from a total of 60 z-slices are shown on this map. This antennal lobe contained 82 glomeruli in total. 64 GFP-labeled glomeruli were annotated: 56 classified as invariant glomeruli and 8 classified as variant glomeruli. Two of the *orco* (+) invariant glomeruli (CD5 and CD6 found between −25 to −31 μm) are not shown in the presented slices. *orco* (+) neurons innervate all of the anterior glomerular groups namely, the Antero-Dorsal (AD), Antero-Lateral (AL), Antero-Medial (AM), Antero-Central (AC) and the Ventro-Central (VC) groups. The posterior groups innervated by *orco* (+) neurons include the Postero-Medial (PM), Postero-Central (PC), Postero-Lateral (PL), Centro-Dorsal (CD), Postero-Ventral (PV), Dorsal (D) and Medio-Dorsal (MD) groups. Glomeruli CD2, CD3, PL3, PL4, PC4, PM4 are putatively co-innervated by *orco* (+) and *IR8a* (+) neurons. 18 other spatially invariant glomeruli without innervation by *orco* (+) neurons are annotated on this map. 8 *orco* (+) variant glomeruli were also annotated on this lobe: AL5, AD6, CD8, D6, VC11, D7 which are shown, and CD7 (−15 to −22 μm) and PM6 (−25 to −30 μm) which are not shown. Scale bar: 10 μm.

**Figure 2—figure supplement 3.**
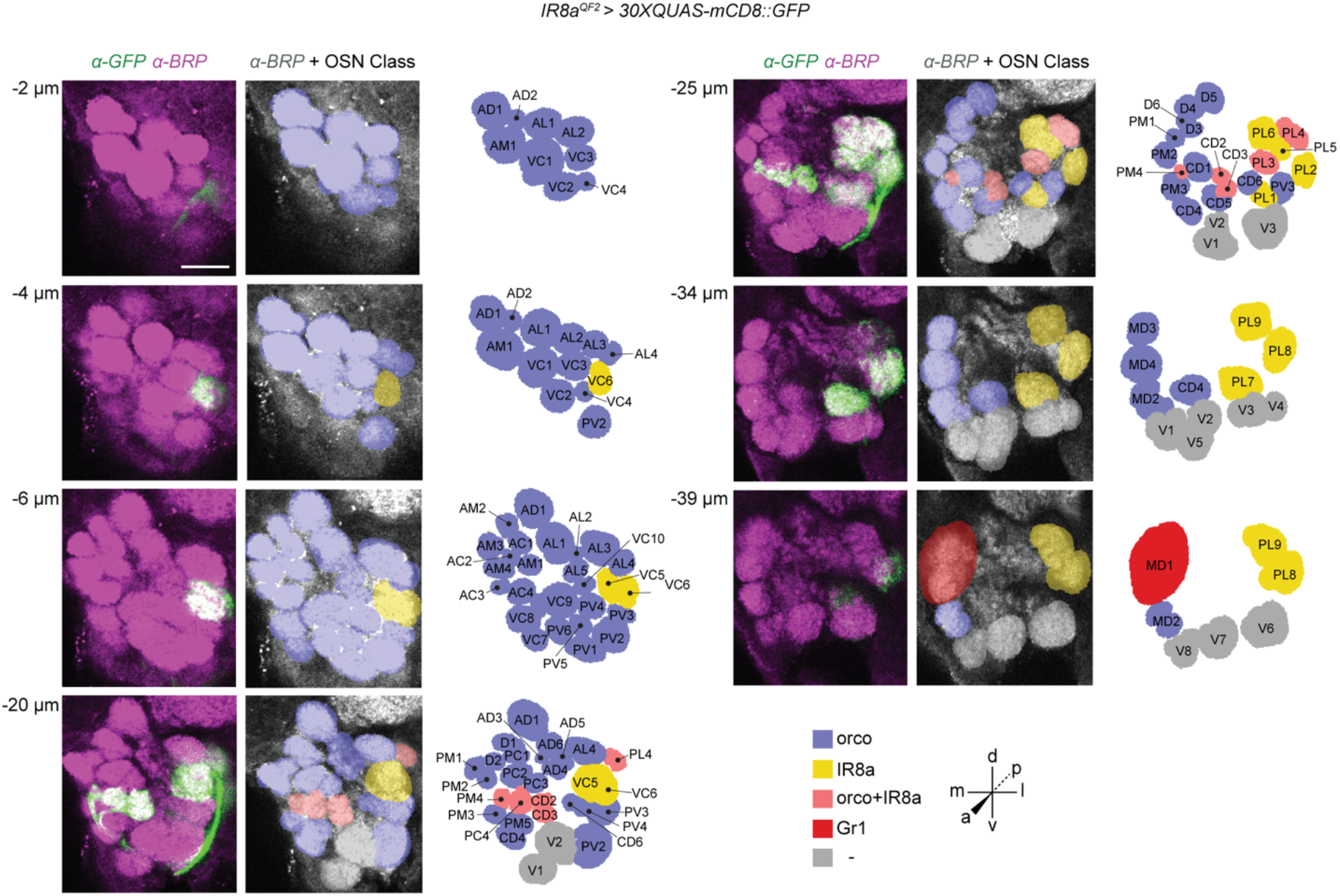
*IR8a* receptor to glomerulus 2D map. Representative confocal stack from the left antennal lobe of an adult female *IR8a^QF2^ > 30XQUAS-mCD8::GFP* mosquito. 7 representative focal planes from a total of 44 z-stacks are shown on this map. This antennal lobe contained 79 glomeruli in total. 15 GFP-labeled glomeruli were annotated that are classified as being spatially invariant: 2 glomeruli of the anteriorly positioned VC group (VC5 and VC6), 9 glomeruli of the PL group (PL1-9), 2 CD glomeruli (CD2 and CD3), 1 PC group glomerulus (PC4) and 1 PM group glomerulus (PM4). Glomeruli CD2, CD3, PL3, PL4, PC4, PM4 are putatively co-innervated by *orco* (+) and *IR8a* (+) neurons. 59 other spatially invariant glomeruli without innervation by *IR8a* (+) neurons are annotated on this map. 5 variant glomeruli were also annotated on this lobe: AL5, AD6, D6 which are shown, and VC11 (−14 to −18 μm) and VC12 (−14 to −19 μm) which are not shown. Scale bar: 10 μm.

**Figure 2—figure supplement 4.**
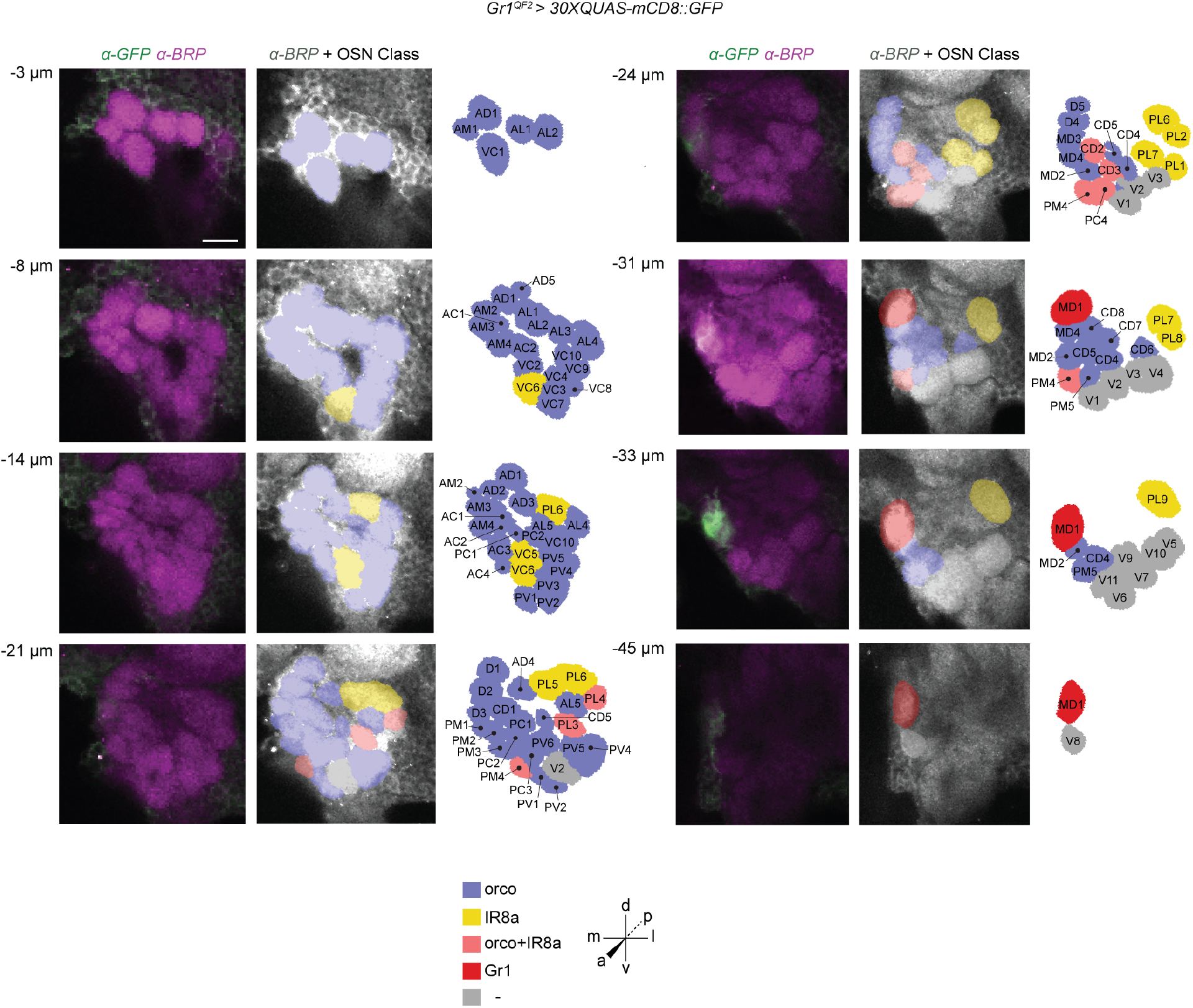
*Gr1* receptor to glomerulus 2D map. Representative confocal stack from the left antennal lobe of an adult female *Gr1^QF2^ > 30XQUAS-mCD8::GFP* mosquito. 8 representative focal planes from a total of 50 z-stacks are shown on this map. This antennal lobe contained 81 glomeruli in total. 1 GFP-labeled glomerulus was annotated. *Gr1* (+) neurons project to the spatially invariant glomerulus MD1. 73 other spatially invariant glomeruli without innervation by *Gr1* (+) neurons are annotated on this map. 7 variant glomeruli were also annotated on this lobe: V9, V10, V11, CD7, CD8 and AL5 which are shown, and V12 (−38 μm) which is not shown. Scale bar: 10 μm.

**Figure 2—figure supplement 5.**
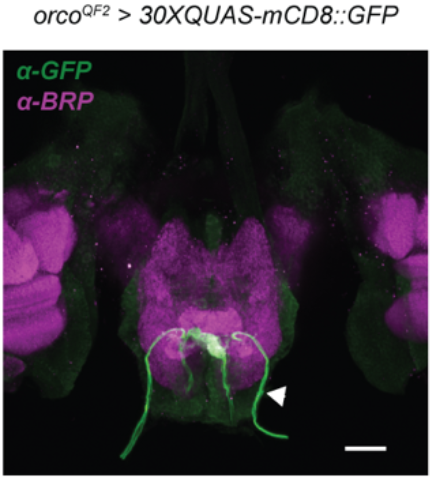
SEZ innervation of *orco* (+) neurons. Posterior view of the brain of an adult female *orco^QF2^ > 30XQUAS-mCD8::GFP* mosquito. *orco* (+) neurons from the labella project via the labial nerve (arrow) and terminate in the gustatory center of the mosquito brain, the subesophageal zone (SEZ). In *Anopheles gambiae,* these terminal neuropil clusters were well separated and named SEZ glomeruli (Riabinina et al., 2016). In *Aedes aegypti* the boundaries of these clusters overlap and appear to be smaller in size. Scale bar: 40 μm.

**Figure 4—figure supplement 1.**
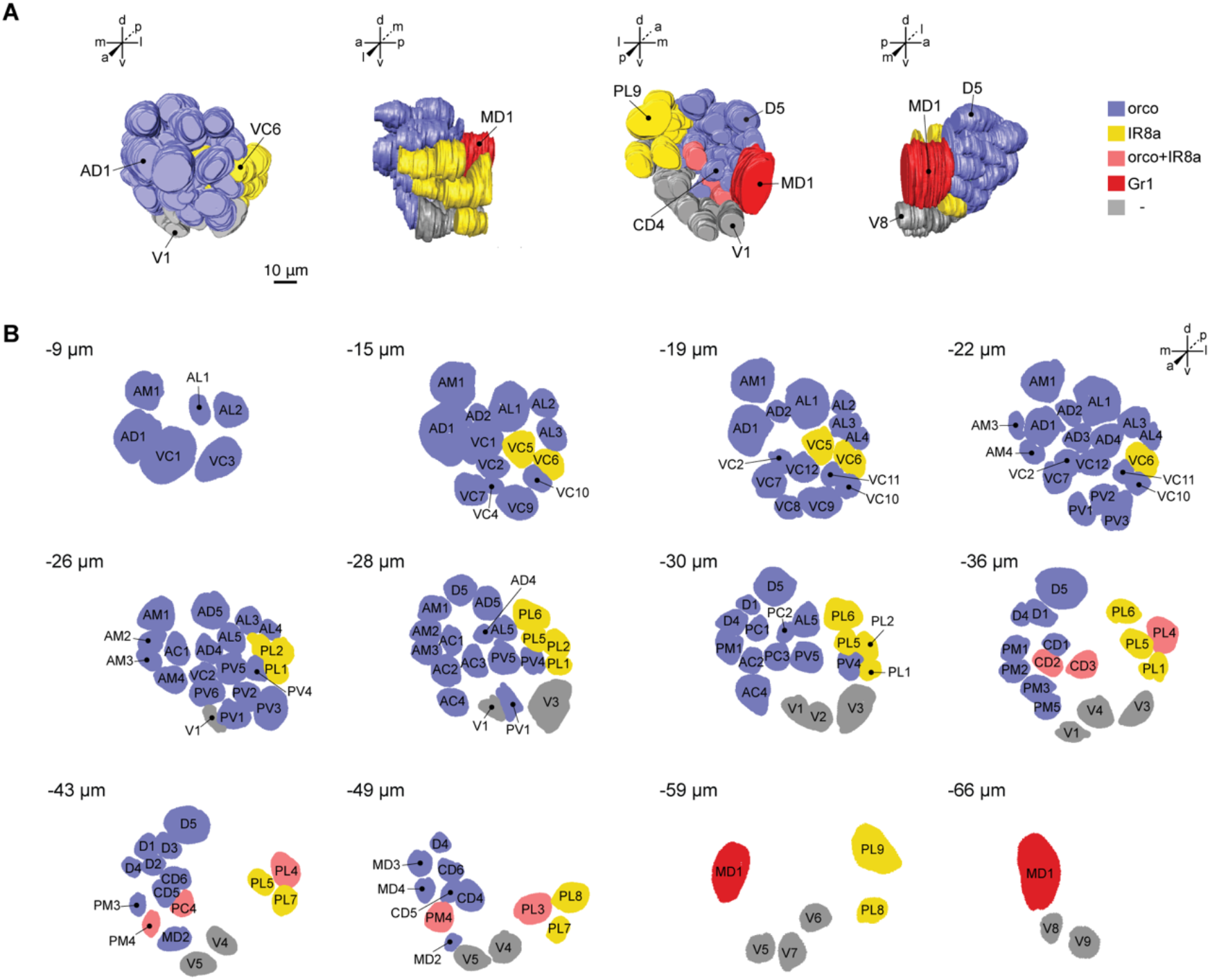
Phalloidin-stained LVPib12 female antennal lobe model used as a reference map for CaMPARI2 activity dependent labeling. **(A)** 3D model generated from a female LVib12 left antennal lobe stained with Alexa Fluor 647-phalloidin and imaged at 63X magnification from anterior, lateral, posterior and medial perspectives. Glomeruli are color coded according to chemoreceptor class. Landmark glomeruli are highlighted on the model. **(B)** 2D reference map for spatial registration of CaMPARI labeling. This antennal lobe contained 82 glomeruli in total. All 74 invariant glomeruli are shown. 8 variant glomeruli were also annotated on this map: V9, VC11, VC12 and AL5 are shown, and CD7, PC5, PV7 and PM6 are not shown and found in intervening slices. Scale bar: 10 μm.

**Figure 5—figure supplement 1.**
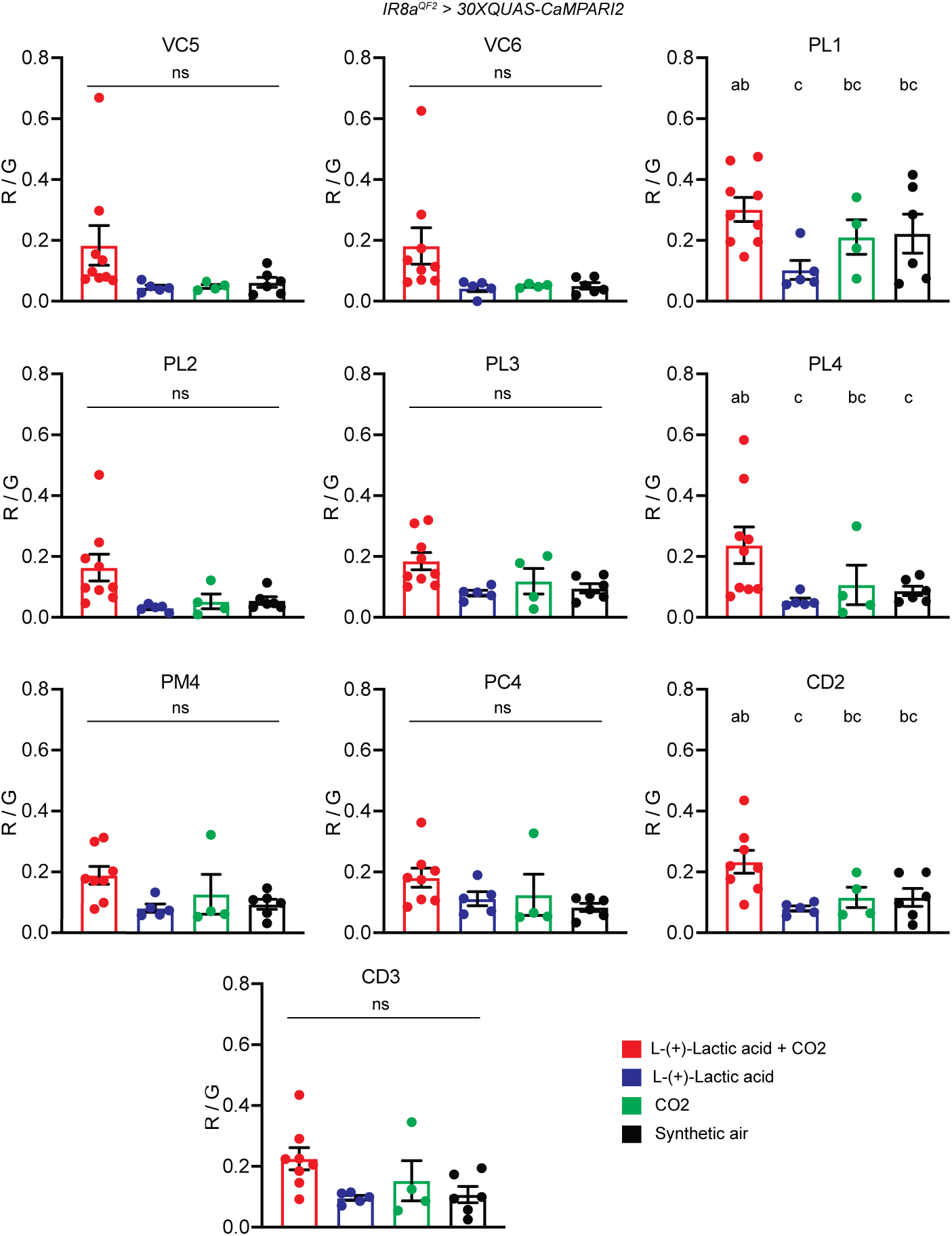
CaMPARI2 photoconversion in supplemental *IR8a* (+) glomeruli screened in response to co-stimulation with L-(+)-lactic acid and CO_2_. Comparisons of CaMPARI2 photoconversion revealed no significant differences in glomerular photoconversion in VC5, VC6, PL2, PL3, PM4 and CD3 glomeruli with any of the four odorant stimuli (L-(+)-lactic acid + CO_2_; L-(+)-lactic acid alone, CO_2_ alone; and synthetic air). Of the remaining glomeruli, PL4, PL5, PL6 and CD2 glomeruli exhibited photoconversion ratios where the ‘L-(+)-lactic acid + CO_2_’ stimulus had mean R/G values that were significantly elevated relative to all or some of the other stimuli. For example, the PL4 glomerulus had mean photoconversion values for ‘L-(+)-lactic acid +CO_2_ vs Synthetic Air’, *P* = 0.0320; and ‘L-(+)- lactic acid + CO_2_ vs L-(+)-lactic acid’, *P* = 0.0094; that were significantly different. For the CD2 glomerulus, differences between mean R/G values for ‘L-(+)-lactic acid + CO_2_ vs L-(+)-lactic acid’ also reached statistical significance (*P* = 0.0481). Tukey’s Multiple Comparison Test, *n* = 4-9 brains per stimulus, mean R/G values ± s.e.m. plotted.

**Table S1.**
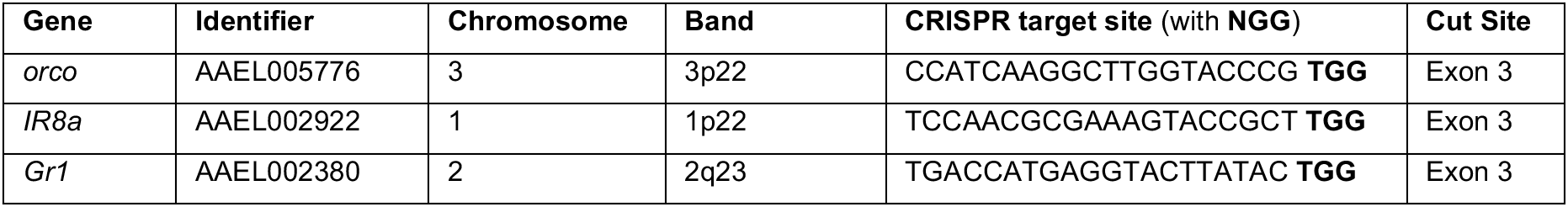
CRISPR target sites and locations.

**Table S2.**
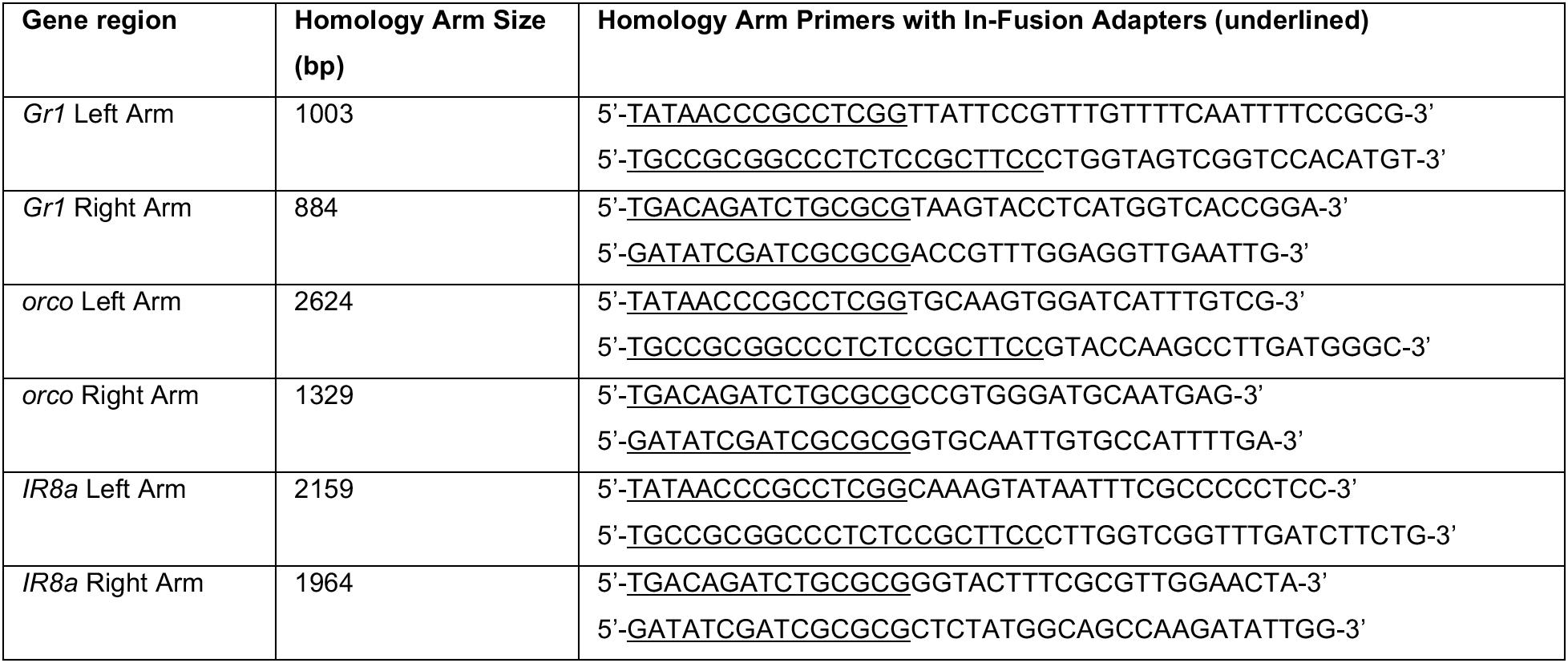
Homology arms for *T2A-QF2* donor constructs.

**Table S3.** Template materials for constructs.

| Plasmid | Element | Source | Addgene # |
| --- | --- | --- | --- |
| pBB | 3xP3-ECFP-SV40 | pBAC-ECFP-15xQUAS-TATA-SV40 | 104875 |
|  | T2A-QF2-hsp70-loxP | pHACK-QF2 | 80274 |
|  | 3xP3 promoter | pBAC-ECFP-15xQUAS-TATA-SV40 | 104875 |
|  | DsRed2-SV40 | pMOS-3xP3-DsRed | n/a |
| All reporter constructs |  |  |  |
|  | Mos1 and ECFP marker | pMos 3xP3-ECFPaf} | n/a |
|  | 15xQUAS promoter | pBAC-ECFP-15xQUAS-TATA-SV40 | 104875 |
|  | Syn21 enhancer | pJFRC81-10XUAS-IVS-Syn21-GFP-p10 | 36432 |
|  | p10 terminator | pJFRC81-10XUAS-IVS-Syn21-GFP-p10 | 36432 |
| pMosECFP-15XQUAS-mCD8::GFP | mCD8:GFP | pQUASp-mCD8::GFP | 46163 |
| pMos-loxP-ECFP-loxP-15XQUAS-CaMPARI2 | CaMPARI2 | pAAV-hsyn1-CaMPARI2 | gift of E. Schreiter |
| pMosEYFP-exu-Cre |  |  |  |
|  | Mos1 | pMOS-3xP3-dsRed | n/a |
|  | *Polyubiquitin* promoter | pSL1180-HR-PUbECFP | 47917 |
|  | EYFP | pBAC-YFP-QF2-hsp70 | gift of C. Potter |
|  | *exuperantia* promoter | AAEL010097-Cas9 mosquitoes | gift of O. Akbari |
|  | Cre recombinase | pENTR L1-vas2-Cre-L4 | 62301 |
|  | p10 terminator | pJFRC81-10XUAS-IVS-Syn21-GFP-p10 | 36432 |

**Table S4.**
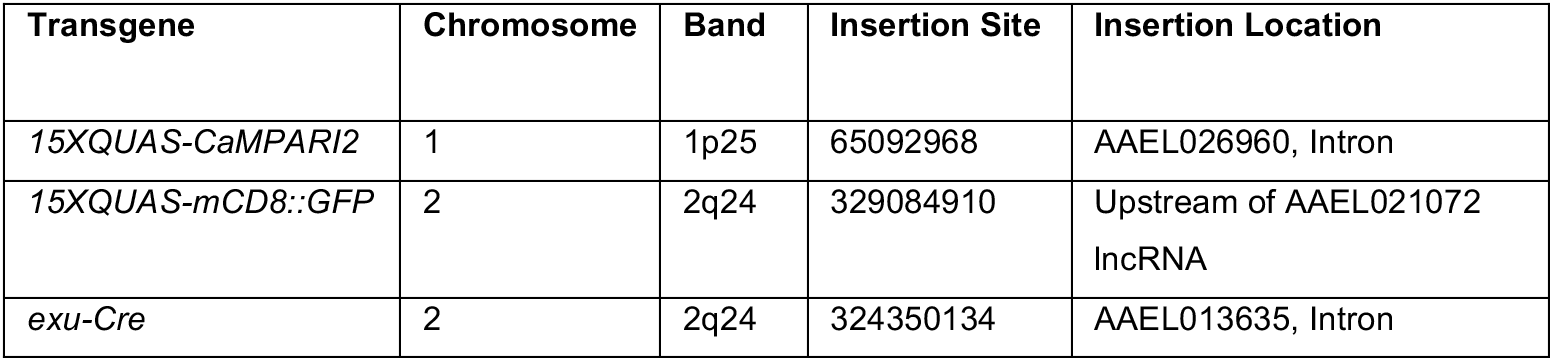
Genomic integration sites of *Mos1 mariner* transgenes.

| Transgene | Chromosome | Band | Insertion Site | Insertion Location |
| --- | --- | --- | --- | --- |
| *15XQUAS-CaMPARI2* | 1 | 1p25 | 65092968 | AAEL026960, Intron |
| *15XQUAS-mCD8::GFP* | 2 | 2q24 | 329084910 | Upstream of AAEL021072<br>lncRNA |
| *exu-Cre* | 2 | 2q24 | 324350134 | AAEL013635, Intron |

## Notes

### Competing Interest Statement

The authors have declared no competing interest.

### Summary of Updates

Manuscript has been revised to include detailed schematics of: (i) CRISPR-Cas9 mediated T2A-in frame fusion strategy to gain genetic access to targeted subsets of Aedes aegypti olfactory sensory neurons, and; (ii) a summary of chemoreceptor innervation and annotation of Aedes aegypti antennal lobe glomeruli. Text in the introduction and discussion has been updated to discuss our results within the context of complementary studies focusing on the neuroanatomical and functional characterization of the Aedes aegypti antennal lobe.

